# Neuroglia Infection by Rabies Virus after Anterograde Virus Spread in Peripheral Neurons

**DOI:** 10.1101/2020.09.20.305078

**Authors:** Madlin Potratz, Luca M. Zaeck, Carlotta Weigel, Antonia Klein, Conrad M. Freuling, Thomas Müller, Stefan Finke

**Affiliations:** Friedrich-Loeffler-Institut (FLI), Federal Research Institute for Animal Health, Institute of Molecular Virology and Cell Biology, 17493 Greifswald-Insel Riems, Germany

**Keywords:** rabies pathology, tissue optical clearing, 3D tissue imaging, light sheet microscopy, neuroglia infection, Schwann cell

## Abstract

The highly neurotropic rabies virus (RABV) enters peripheral neurons at axon termini and requires long distance axonal transport and trans-synaptic spread between neurons for the infection of the central nervous system (CNS). Whereas laboratory strains are almost exclusively detected in neurons, recent 3D imaging of field RABV-infected brains revealed an remarkably high proportion of infected astroglia, indicating that in contrast to attenuated lab strains highly virulent field viruses are able to suppress astrocyte mediated innate immune responses and virus elimination pathways. While fundamental for CNS invasion, *in vivo* field RABV spread and tropism in peripheral tissues is understudied. Here, we used three-dimensional light sheet and confocal laser scanning microscopy to investigate the *in vivo* distribution patterns of a field RABV clone in cleared high-volume tissue samples after infection via a natural (intramuscular; hind leg) and an artificial (intracranial) inoculation route. Immunostaining of virus and host markers provided a comprehensive overview of RABV infection in the CNS and peripheral nerves after centripetal and centrifugal virus spread. Importantly, we identified non-neuronal,axon-ensheathing neuroglia (Schwann cells, SCs) in peripheral nerves of the hind leg and facial regions as a target cell population of field RABV. This suggest that virus release from axons and infected SCs is part of the RABV *in vivo* cycle and may affect RABV-related demyelination of peripheral neurons and local innate immune responses. Detection of RABV in axon surrounding myelinating SCs after i.c. infection further provided evidence for anterograde spread of RABV, highlighting that RABV axonal transport and spread of infectious virus in peripheral nerves is not exclusively retrograde. Our data support a new model in which, comparable to CNS neuroglia, SC infection in peripheral nerves suppresses glia-mediated innate immunity and delays antiviral host responses required for successful transport from the peripheral infection sites to the brain.

## Introduction

Rabies is estimated to cause 59.000 human deaths annually in over 150 countries, with 95% of cases occurring in Africa and Asia (reviewed in [26,21]). The disease is caused by a plethora of highly neurotropic, non-segmented, single-stranded RNA viruses of negative polarity belonging to the family *Rhabdoviridae* of the order *Mononegavirales,* of which the rabies virus (RABV) is the type species of the Lyssavirus genus [2]. RABV and other lyssaviruses enter the nervous system at the synapses of peripheral neurons. After receptor-mediated endocytosis, the virions remain in endosomal vesicles and hijack the retrograde axonal transport machinery [33,23,59] to reach the neuron soma, where virus ribonucleoproteins (RNPs) are released into the cytoplasm [47] for virus gene expression and replication. Newly formed virus is then transported to post-synaptic membranes, followed by trans-synaptic spread to connected neurons [12]. Prolonged replication of pathogenic RABV in the brain eventually leads to the onset of a progressive rabies encephalitis, invariably resulting in the fatal disease progression. In animals suffering from RABV encephalitis, the virus can be detected in multiple nervous or innervated tissues including hind leg, spinal cord, brain, face, and salivary glands [5,15]. For the transmission of infectious virus to new hosts via virus-containing saliva, centrifugal spread of RABV in the peripheral nervous system (PNS) is required [18]. The secretion of RABV-containing saliva by salivary glands is commonly accepted as the main source of virus shedding and virus transmission to uninfected animals via bites (reviewed in [21,16]). Detection of RABV in sensory neurons of the olfactory epithelium and other tissues during late stages of infection indicates that, at least in these late phases of disease, the axonal transport is not exclusively retrograde (reviewed in [16]). In fact, studies have shown that RABV enters the anterograde axonal transport pathway in cultivated primary rat dorsal root ganglion (DRG) neurons [6] and in mice [70]. However, the consequences of anterograde transport of RABV is still controversially discussed, even though infection through intradermal sensory neurons represents a likely route of entry for lyssaviruses as a result from bat bites [7].

Because of their trans-synaptic spread, RABVs constitute ideal viruses to map neuronal connections [67,66] as even attenuated RABVs enter axonal transport pathways [33]. More neuroinvasive yet still lab-adapted strains like CVS-11 have been used as polysynaptic neuron tracers and the resultant insights have subsequently been used to explain *in vivo* RABV spread and pathogenesis [61]. However, already between different CVS strains, substantial differences in pathogenicity exist [39], raising the question whether knowledge from established lab-adapted strains can be extrapolated to explain actual field RABV infection and pathogenesis.

The advent of efficient full-genome cloning techniques opened novel avenues for the analysis of virus variability in combination with phenotypical characterization of recombinant field viruses at a clonal level [43]. For example, a recent comparison of field and lab RABV clones in mice revealed that field viruses not only showed a higher virulence compared to lab strains such as CVS-11 but also substantially differ in their ability to establish infection in non-neuronal astroglia in the central nervous system (CNS) [48]. In contrast to attenuated lab-adapted RABV, which is supposed to be eliminated from infected glial cells (astrocytes) by type I interferon-induced immune reactions [46], it has been hypothesized that rapid onset of field virus gene expression in non-neuronal astrocytes blocks inhibitory innate immune activation and responses [48]. This may not only affect RABV replication in glial cells directly but also in surrounding neurons via paracrine signalling. Thus, this mechanism may be crucial to RABV spread and pathogenesis *in vivo*. Apart from astrocytes, *in vivo* infection of non-neuronal muscle, epidermal, and epithelial cells have been described [12,5,13,42]. In foxes, which had been orally immunized with live-attenuated RABV, infected cells can be detected in the epithelium of lymphoid tonsils [63,56]. *In vitro*, even immature DCs or monocytes can be infected [35]. Overall, knowledge on the broad spectrum infection pattern of RABV *in vivo,* particularly of field RABV strains, is still incomplete and the identification of PNS and non-neuronal cells involved in RABV spread and pathogenesis has been given little attention.

Therefore, by using a highly virulent field RABV clone of dog origin (rRABV Dog) [43], we set out to systematically and comprehensively elucidate the *in vivo* spread and cell tropism of a field RABV strain. So far, technical limitations in conventional histology to visualize infection processes in complex tissues in large three-dimensional (3D) volumes at a mesoscopic scale and high resolution hindered a comprehensive overview of RABV cell tropism, *in vivo* spread, and pathogenesis. Immunofluorescence-compatible tissue clearing iDISCO and uDISCO (immunolabeling-enabled and ultimate three-dimensional imaging of solvent-cleared organs, respectively) protocols [45,49] recently allowed us to perform high-resolutions 3D immunofluorescence analyses of thick RABV-infected tissues slices [69,48].

Here, we investigated the *in vivo* spread of a highly virulent field RABV clone (rRABV Dog) through the CNS and PNS after intramuscular (i.m.) and intracranial (i.c.) inoculation. Using 3D immunofluorescence imaging of solvent-cleared tissues by light sheet and confocal laser scanning microscopy, we identified RABV infections in hind leg nerves, spinal cord, brain, and nerves of the facial region. These data provide a broad overview of field RABV distribution in the clinical phase of rabies encephalitis. Notably, we were able to identify Schwann cells (SCs) as a so far unrecognized non-neuronal RABV target cell population, which may be involved in regulating RABV infection and, thus, pathogenesis in peripheral nerve tissues. Moreover, infection of SCs after centrifugal RABV spread provides evidence for anterograde axonal transport and infectious virus release at distal axon membrane sites of peripheral neurons.

## Materials and Methods

### Viruses

The recombinant field virus clone rRABV Dog [43] was derived from an isolate originating from an infected dog from Azerbaijan, which was used in a challenge study [64]. RABV Bat (FLI virus archive ID N 13240) was isolated from an insectivorous North American bat.

### Animal trials

Six-to eight-week-old BALB/c mice (Charles-River, Germany) were i.m.-infected with decreasing doses of rRABV Dog. Four groups of six animals each were anesthetized with isoflurane and infected with 3×10^0^, 3×10^1^, 3×10^2^ and 3×10^3^ TCID_50_/mouse by injection of 30 μL of virus suspension in the femoral hind leg muscle. Additionally, three mice were i.c.-infected at a dose of 1×10^2^ TCID_50_/mouse. Body weight and clinical scores from one to five (1: ruffled fur, hunched back; 2: slowed movement, circular motions, weight loss > 15%; 3: tremors, shaky movement, seizures, weight loss > 20%; 4: paralysis, weight loss > 25%; 5: coma/death) were determined daily until day 21 post-infection or day of euthanasia. After reaching a clinical score of two, the animals were anaesthetized with isoflurane and euthanized by decapitation. Samples were taken, fixed with 4% paraformaldehyde (PFA) for one week, and stored for further processing. All remaining animals were euthanized at 21 dpi. The animal approaches were evaluated by the responsible animal care, use, and ethics committee of the State Office for Agriculture, Food Safety, and Fishery in Mecklenburg-Western Pomerania (LALFF M-V) and gained approval with permission 7221.3-2-047/19). For RABV Bat, archived tissue samples (hind legs, spinal columns, and heads) from previous virus characterization experiments were used (LALFF M-V permission 7221.3-2.1-001/18).

### Detection of RABV RNA in oral swabs and salivary glands via RT-PCR

Salivary gland samples were homogenized in 1 ml cell culture medium in the UPHO Ultimate Homogenizer (Geneye, Hong Kong) using a 3 mm steal bead. Oral swab samples were transferred into 1 ml of culture medium. Total RNA from oropharyngeal swabs as well as from salivary gland tissues were extracted using the NucleoMagVet kit (Macherey&Nagel, Germany) according to manufacturer’s instructions in a KingFisher/BioSprint 96 magnetic particle processor (Qiagen, Germany). RABV RNA was detected by the R14-assay, a RT-qPCR targeting the N-gene [27]. The RT-qPCR reaction was prepared using the AgPath-ID-One-Step RT-PCR kit (Thermo Fisher Scientific, USA) adjusted to a volume of 12.5 μl [19]. RT-qPCRs were performed on a BioRad real-time CFX96 detection system (Bio-Rad, USA).

### Antibodies and dyes

To detect RABV in the tissues, a polyclonal rabbit serum against recombinant RABV P protein (P160-5; 1:3,000 in PTwH [0.2% Tween 20 in PBS with 10 μg/mL heparin]) was used [44]. The following first antibodies and dyes were obtained from the respective suppliers: chicken anti-NEFM (Thermo Fisher, USA; #PA1-16758, RRID:AB_2282551; 1:1,000 in PTwH), guinea pig anti-NEFM (Synaptic Systems, Germany; #171204, RRID:AB_2619872; 1:400 in PTwH), chicken anti-MBP (Thermo Fisher; #PA1-10008, RRID:AB_1077024; 1:500 in PTwH), TO-PRO™-3 iodide (Thermo Fisher; #T3605; 1:1,000 in PTwH.) As secondary antibodies donkey anti-rabbit Alexa Fluor® 568 (Thermo Fisher; #A10042, RRID:AB_2534017), donkey anti-chicken Alexa Fluor® 488 (Dianova, Germany; #703-545-155, RRID:AB_2340375), donkey anti-guinea pig Alexa Fluor® 647 (Dianova; #706-605-148, RRID:AB_10895029) were used, each at dilution of 1:500 in PTwH.

### iDISCO-based immunostaining and uDISCO-based clearing of tissue samples

Staining and clearing protocols were adapted from earlier reports [45,49] and were performed as described previously [69,48]. Hind legs were skinned prior to fixation. Prior to the slicing, PFA-fixed hard tissues, which included hind legs, spinal columns, and heads, were decalcified with 20% ethylenediaminetetraacetic acid (EDTA) [w/v] in H_2_O_dd_ (pH was adjusted to 7.0) for four days at 37 °C. The EDTA solution was exchanged daily. Afterwards the decalcified tissues were cut into 1 mm thick sections using a scalpel. For brains, as soft tissues, a vibratome (VT1200S, Leica Biosystems, Germany) was used. After treatment with increasing methanol concentrations in distilled water (20%, 40%, 60%, 80%, and twice 100%) and bleaching with 5% H_2_O_2_ in pure methanol at 4 °C, the samples were rehydrated with decreasing concentrations of methanol in distilled water (80%, 60%, 40%, and 20%). After washing with PBS and 0.2% Triton X-100 in PBS, the samples were permeabilized for 2 days at 37 °C in 0.2% Triton X-100/20% DMSO/0.3 M glycine in PBS. After subsequent blocking with 6% donkey serum/0.2% Triton X-100/10% DMSO in PBS at 37 °C for another 48 h, primary antibodies in PTwH were added for 5 days with refreshment of the antibody solution after 2.5 days. The samples were washed with PTwH four times until the next day, when secondary antibodies in PTwH were added for 5 days with refreshment of the antibody solution after 2.5 days. After further washing for four times until the next day, the tissues were dehydrated by a series of *tert*-butanol (TBA) solutions (from 30% to 96% in distilled water), each step for 2 h (except over night incubation with for 80% TBA), final incubation in 100% TBA and subsequent clearing in BABB-D15 with 0.4 vol% DL-α-tocopherol until they were optically transparent (2 – 6 h). BABB-D15 is a combination of benzyl alcohol (BA) and benzyl benzoate (BB) at a ratio of 1:2, which is mixed at a ratio of 15:1 with diphenyl ether (DPE). The samples were either analyzed directly by light sheet microscopy or were transferred to 3D-printed imaging chambers as described before (Zaeck et al., 2019.)

### Light sheet and confocal laser scanning microscopy

For light sheet microscopy, a LaVision UltraMicroscope II with an Andor Zyla 5.5 sCMOS Camera, an Olympus MVX-10 Zoom Body (magnification range: 0.63 – 6.3x), and an Olympus MVPLAPO 2x objective (NA = 0.5) equipped with a dipping cap was used. Z-stacks were acquired with a light sheet NA of 0.156, a light sheet width of 100%, and a z-step size of 2 μm. LaVision BioTec ImSpector Software (v7.0.127.0) was used for image acquisition.

Confocal laser scanning microscopy was performed with a Leica DMI 6000 TCS SP5 equipped with a HC PL APO CS2 40x water immersion objective (NA = 1.1; Leica, Germany; #15506360), using the Leica Application Suite Advanced Fluorescence software (V. 2.7.3.9723) and a z-step size of 0.5 μm.

### Image processing

For image processing, a Dell Precision 7920 workstation was used (CPU: Intel Xeon Gold 5118, GPU: Nvidia Quadro P5000, RAM: 128 GB 2666 MHz DDR4, SSD: 2 TB). Acquired image stacks were processed using the ImageJ (v1.52h) distribution package Fiji [53,54] or arivis Vision4D (arivis, Germany; v3.2.0). Brightness and contrast were adjusted for each channel and, if necessary, channels were denoised. Either maximum-z-projections or volumetric 3D-projections are shown.

## Results

### Field virus clone rRABV Dog undergoes the complete RABV infection cycle in adult BALC/c mice

Mouse models differ in their age-dependent susceptibility to fixed RABV lab strains and field isolates [11]. To confirm the field virus character of rRABV Dog and its suitability for investigation of the complete RABV *in vivo* cycle from infected muscle at the site of entry to secretion in saliva in adult mice, six-to eight-week-old BALB/c mice were inoculated by the i.m. route (hind leg muscle) with decreasing infectious virus doses. Although mortality was dose-dependent, even very low doses (30 TCID_50_/mouse) of rRABV Dog were sufficient to cause rabies disease and release of infectious virus in nearly all inoculated mice, thus, highlighting the high virulence of this field virus RABV clone. Comparable to field RABV isolates, even at an extremely low infection dose (3 TCID_50_/mouse), the complete *in vivo* cycle had been undergone by animals that succumbed to infection, as shown by detection of viral RNA in both salivary glands and in oral swabs using RT-qPCR (Table 1; Fig. S1).

**Table 1:** Dose-dependent onset of clinical rabies signs after i.m. application of rRABV Dog in the upper hind leg muscle of six-to eight-week old BALB/c mice. Salivary gland and oral swabs were virus RNA positive even at low infection doses. *One oral swab not tested.

| virus dose | euthanized | salivary gland | oral swab |
| --- | --- | --- | --- |
| TCID<sub>50</sub>/mouse | (humane endpoint reached) | virus RNA | virus RNA |
| 3,000 | 6/6 | 6/6 | 5*/6 |
| 300 | 5/6 | 5/6 | 5/6 |
| 30 | 5/6 | 5/6 | 5/6 |
| 3 | 2/6 | 2/6 | 2/6 |

### RABV infects peripheral neuron-associated neuroglia in the hind leg after i.m. inoculation

Immunostaining of solvent-cleared hind leg slices of diseased animals for RABV phosphoprotein (P), neurofilament M (NEFM), and cell nuclei (TO-PRO™-3) was performed for light sheet and confocal laser scanning microscopy analysis (Fig. 1). RABV infection of peripheral nerves innervating the femoral tissue could be demonstrated (Fig. 1 A-C, red). Reconstruction of a 3D femoral tissue volume allowed visualization of the distribution of infected nerve fibres in their spatial environment (Fig. 1B, Video S1). The specificity of the RABV immunofluorescence was confirmed by using non-infected hind leg tissues as negative controls (Fig. S3). At a higher resolution, axons were identified via staining by the neuronal cytoskeleton marker (NEFM) (Fig. 1D,E, green) but, notably, did not co-localize with RABV P (Fig. 1E). As RABV-infected peripheral nerves could be visualized in all serial femoral slices analysed (Fig. 1F-J), our approach proved to be appropriate to reconstruct a 3D model of RABV infection patterns in peripheral tissues.

**Fig. 1.**
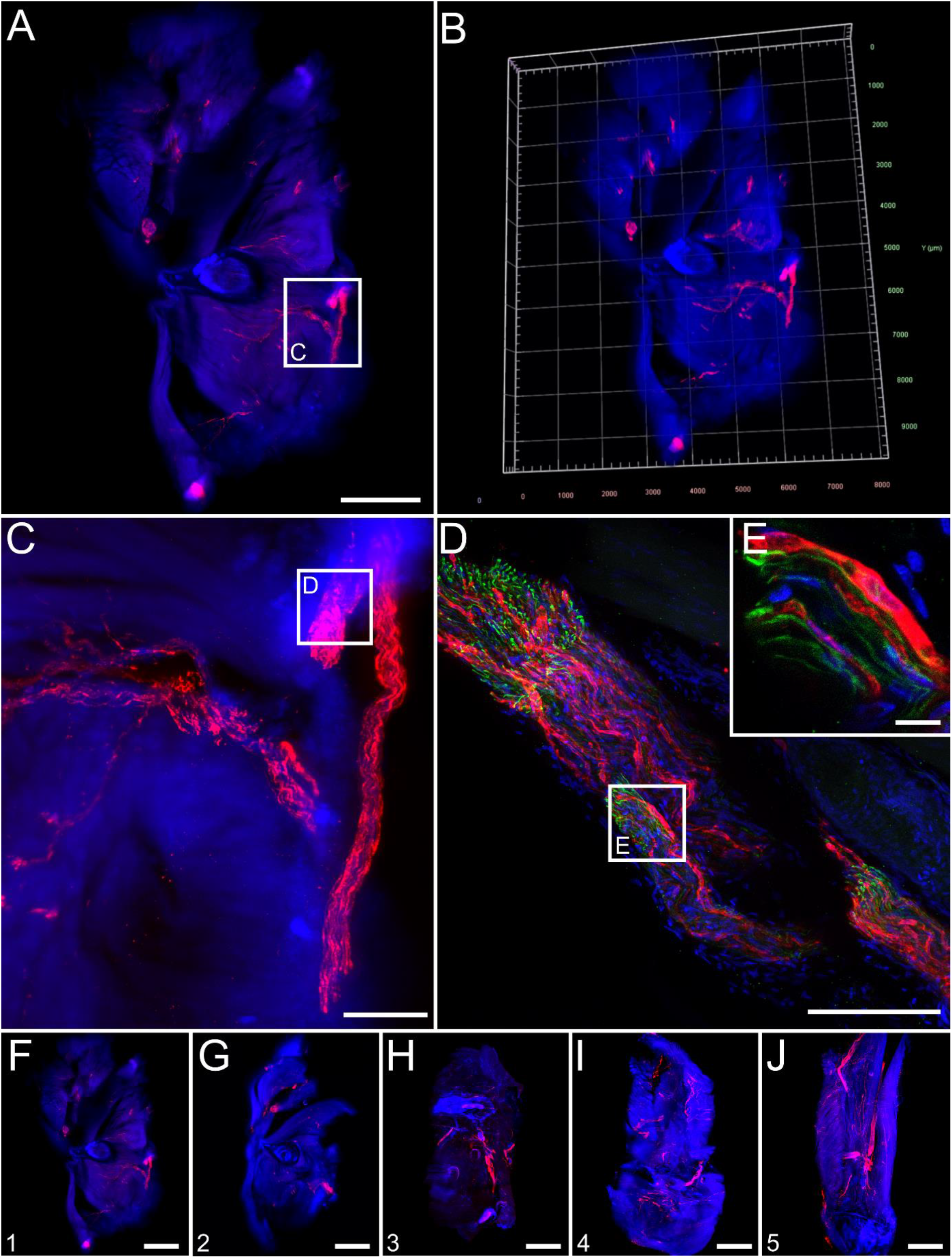
Overview of field RABV infection in mouse hind leg cross sections after i.m. infection (9 days post-inoculation). **(A)** Maximum z-projection of light sheet overview of a hind leg cross section [1.6x magnification; z = 1,156 μm]. Red: RABV P; blue: nuclei. Green fluorescence for NEFM not shown as separation from green autofluorescence was not possible at low resolution. Scale bar: 1,500 μm. **(B)** 3D projection of (A). **(C)** Maximum z-projection of detail (white box in A) with infected nerve [12.6x magnification; z = 574 μm]. Scale bar: 200 μm. **(D)** Maximum z-projection of confocal z-stack of indicated region in (C) [z = 48 μm] of infected nerve with NEFM signal (green). Scale bar: 100 μm. Note: due to the separate image acquisition, the orientation of the shown nerve is different to (C). **(E)** Maximum z-projection of detail from (D) [z = 10 μm]. Scale bar: 10 μm. **(F-J)** Maximum z-projections of serial thick hind leg sections from the upper leg (F) to the knee (J) [1.6x magnification, z = 1,156 μm (F), 1,402 μm (G), 1,172 μm (H), 1,990 μm (I), 1,372 μm (J)]. For reference, see indicated slices in Fig. S2A. For improved intelligibility, only RABV P (red) and nuclei (blue) are shown. Scale bar: 1,500 μm.

Further confocal laser scanning analysis confirmed the lack of co-localization of RABV P with NEFM in axons (Fig. 2A,B). Rather, RABV P specific signals were arranged next to or around the tubular NEFM signals (Fig. 2C,D, arrows; Video S2). Similar spatial arrangement patterns were observed after staining for the Schwann cell (SC) marker myelin basic protein (MBP) (Fig. 2E-K). MBP is known to be part of the myelin sheath of SCs that forms tubular structures around the axons (reviewed in [51]) and accordingly, appeared as hollow tubules that were surrounded by RABV P (Fig. 2 G-K, arrowheads; Video S3). Corroborated by the presence of nuclei in RABV-infected cells (Fig. 2K), this suggests that RABV P accumulated in the cytoplasm of axon-ensheathing SCs. This provides evidence for the infection of and virus gene expression in neuroglia of the PNS in the clinical phase of RABV encephalitis after i.m. infection.

**Fig. 2.**
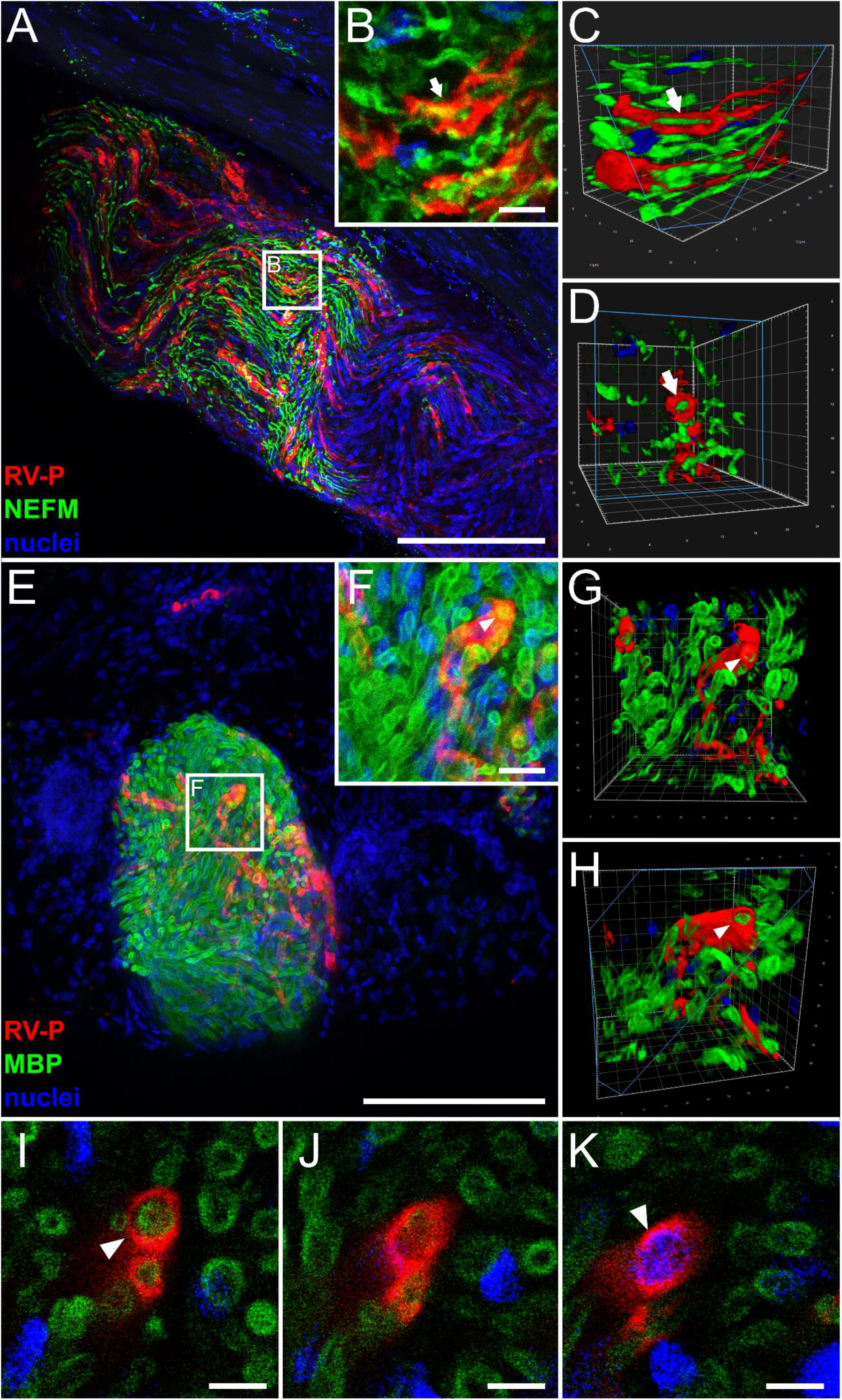
Detection of RABV P in non-neuronal Schwann cells of hind leg tissue after i.m. inoculation (9 days post-inoculation). **(A,B)** Maximum z-projection of high-resolution confocal z-stacks of RABV-infected nerve fiber (see also Fig. 1D) in hind leg sections [z = 45 μm; Scale bar: 100 μm] (A) and detail [z = 15 μm; Scale bar: 10 μm] (B). Red: RABV P; green: NEFM; blue: cell nuclei. **(C,D)** Volumetric 3D projections of (B) with different viewing angles. The light blue box indicates the clipping plane. Arrows: RABV P surrounding axonal NEFM. **(E,F)** Maximum z-projection of a hind leg section (E) [z = 134 μm; Scale bar: 200 μm] and detail (F) [z = 20 μm; Scale bar: 10 μm] stained for RABV P (red), MBP (green) and nuclei (blue). **(G,H)** Volumetric 3D projections of different viewing angles of (F). Arrowhead: RABV-infected Schwann cell **(I-K)** Single slices with RABV P (red), MBP (green) and nuclei (blue). Arrowheads indicate myelin sheath/nucleus surrounded by RABV P. Scale bar: 5 μm.

### Infection of Schwann cells after i.c. inoculation strongly supports anterograde spread of RABV into the periphery

Late phase infection of myelinating SCs in the PNS of i.m.-infected mice could be a result of unspecific virus release from abundantly and long-term RABV-infected myelinated neurons. To investigate, whether RABV also enters the PNS after centrifugal spread from the brain and whether this also leads to infection of myelinating SCs, femoral tissue from i.c.-infected mice was analyzed.

Detection of RABV antigen in the PNS revealed that innervating nerves in femoral tissue can get infected after centrifugal spread of the virus from the brain (Fig. 3A-G). As observed for i.m. inoculations, RABV P fluorescence signals surrounded rather than overlapped with NEFM-positive axons (Fig. 3D,E). Staining for RABV P, MBP, and NEFM (Fig. 3F-K) revealed a three-layered structure with RABV P-specific signals surrounding the MBP-sheath of axons (Fig. 3H-K, Video S4). Infection of myelinated SCs after i.c. inoculation demonstrated that infectious virus was released from peripheral axons, infecting the surrounding axon-associated, non-neuronal cells.

**Fig. 3.**
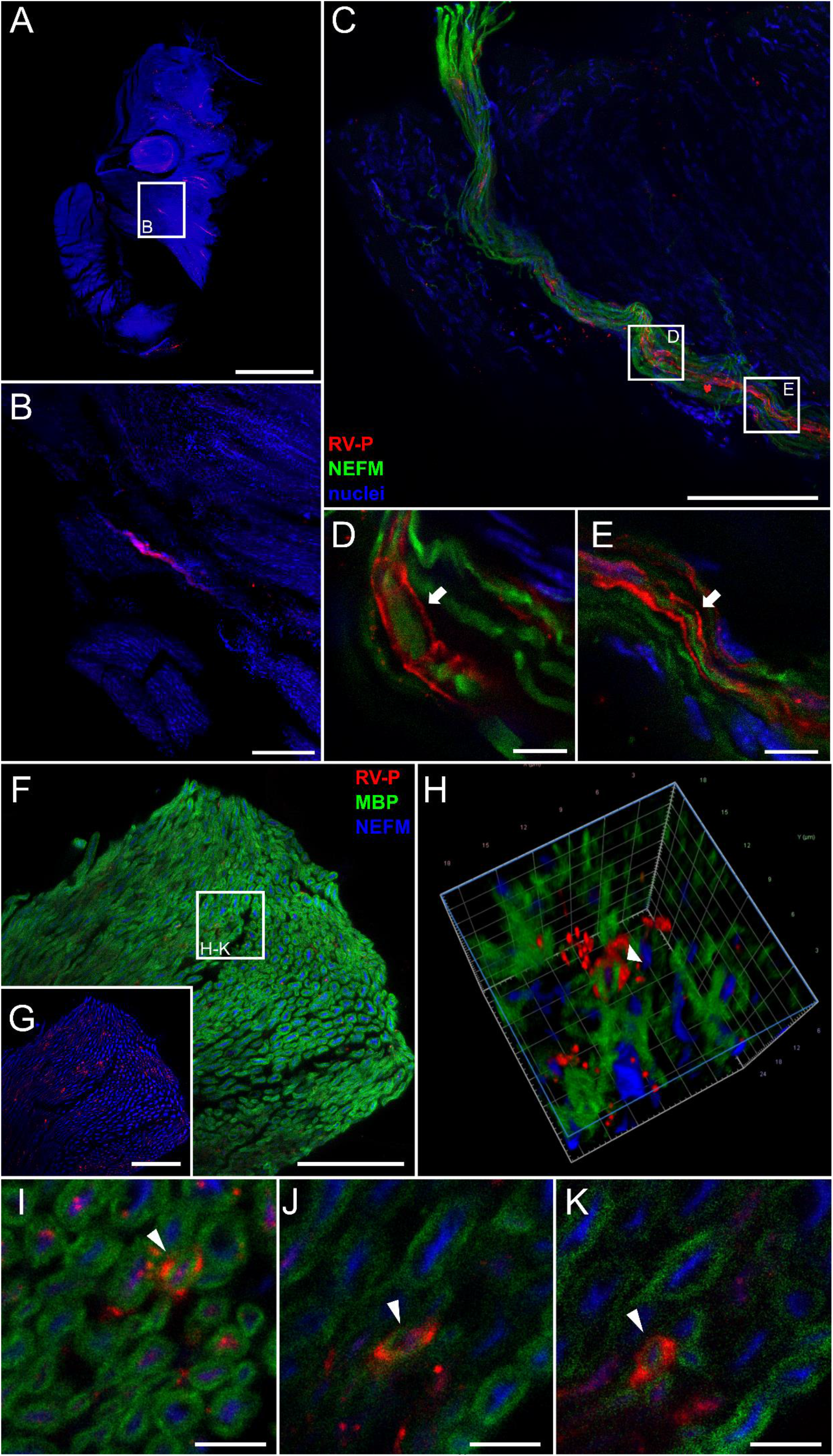
Anterograde virus spread: RABV in Schwann cells of a hind leg nerve after i.c. inoculation. **(A,B)** Maximum z-projection of light sheet overview (A) and detail (B) of hind leg section after staining for RABV P (red), and nuclei (blue). Green fluorescence for NEFM is not shown as separation from green autofluorescence was not possible at low resolution. (A) Magnification of 1.6x [z = 1,944 μm; Scale bar: 1,500 μm] and (B) magnification of 12.6x [z = 58 μm; Scale bar: 200 μm]. **(C)** Maximum z-projection of confocal z-stack [z = 37 μm] of infected nerve fiber from (B), now including NEFM staining (green). Scale bar: 100 μm. **(D,E)** Maximum z-projection of details from (C) (see white boxes) [z = 5 μm (D,E)]. Scale bar: 10 μm. **(F,G)** Maximum z-projection of hind leg nerve section stained for RABV P (red), MBP (green) and NEFM (blue) [z = 15 μm]. To visualize low amounts of viral antigen, the MBP channel is excluded in (G). Scale bar: 50 μm. **(H)** 3D projection of detail from (F) (see white box). **(I-K)** Single planes from detail of (F). The arrowhead indicates an RABV-infected cell. Scale bar: 5 μm.

### RABV exhibits a strong, infection route-independent infection of the spinal cord

To assess the degree of infection of the spinal cord by field virus clone rRABV Dog after different routes of infection, spinal column slices (Fig. S2C) of i.m.- and i.c.-infected mice were systematically analyzed. Independent of the inoculation route, multiple neurons in the spinal cord and neurons projecting out of the same were positive for RABV P (Fig. 4).

**Fig. 4.**
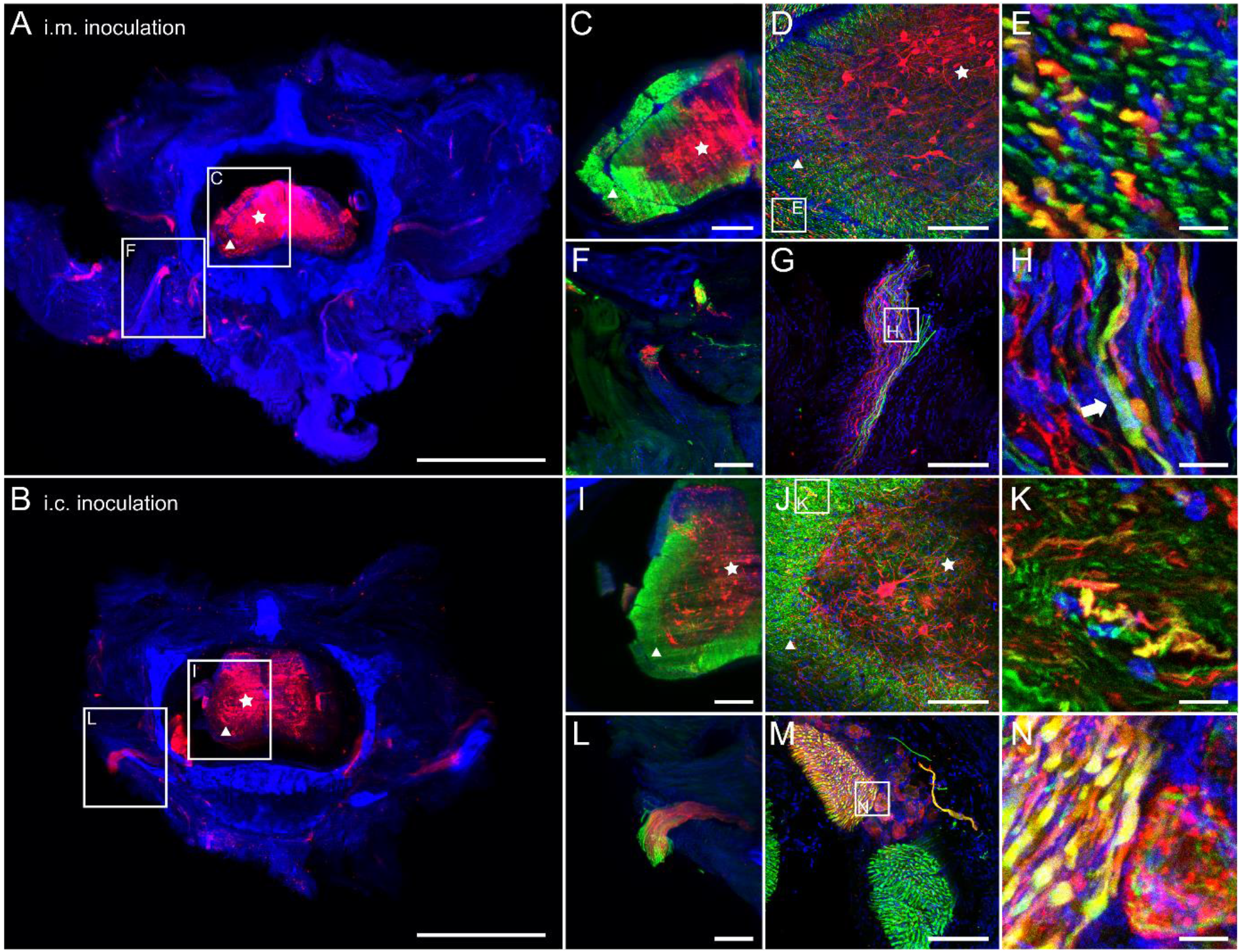
Spinal column infection after i.m. and i.c. infection. **(A,B)** Maximum z-projections of light sheet overviews of cross sections of the spinal column after i.m. (A) and i.c. (B) infection [2.0x magnification; z = 824 μm (A), 1,480 μm (B)]. Red: RABV P; blue: nuclei. Green fluorescence for NEFM not shown as separation from green autofluorescence was not possible at low resolution. Stars: grey matter, triangles: white matter. Scale bar: 1,500 μm. **(C)** Maximum z-projection of detail from (A), including NEFM signals (green). Scale bar: 200 μm. **(D,E)** Maximum z-projection of confocal z-stacks of infected spinal cord (D) [z = 31 μm; Scale bar: 100 μm] and detail (E) [z = 10 μm; scale bar: 10 μm]. **(F)** Maximum z-projection of detail from (A) with neurons projecting from the spinal cord. Scale bar: 200 μm. **(G,H)** Maximum z-projection of confocal z-stacks (G) [z = 36 μm, Scale bar: 100 μm] and detail (H) [z = 10 μm; Scale bar: 10 μm] of projecting nerves from the spinal cord. **(I)** Maximum z-projection of detail from (B), including NEFM signals (green). Scale bar: 200 μm. **(J,K)** Maximum z-projection of confocal z-stacks of infected spinal cord (J) [z = 61 μm; Scale bar: 100 μm] and detail (K) [z = 20 μm; scale bar: 10 μm]. **(L)** Maximum z-projection of detail from (B), including NEFM signals (green). Scale bar: 200 μm. **(M,N)** Maximum z-projection of confocal z-stacks of projecting nerve (M) [z = 41 μm; Scale bar: 100 μm] and detail (N) [z = 20 μm; scale bar: 10 μm].

Higher magnification image stacks of the spinal cord revealed RABV P staining to be primarily localized in the grey matter, surrounded by NEFM-positive white matter (Fig. 4C,I, star and triangle, respectively). Indeed, white and grey matter (Fig. 4D,J, star and triangle, respectively) revealed multiple NEFM-positive axons of which some were infected (Fig. 4E,K). In contrast to the peripheral femoral axons, where the RABV P staining almost exclusively surrounded the tubular NEFM structures (Fig. 2C,D and 3D,E), here, RABV P- and NEFM-specific signals perfectly co-localized (Fig. E,K), suggesting that the axons contained high levels of RABV P protein. Peripheral nerves projecting from the spinal cord also revealed double positive axons (Fig. 4F-H, L-N). However, also RABV P not overlapping with NEFM was observed (Fig. 4H, arrow). In some projecting axon bundles (Fig. 4L), almost all axons were RABV-positive as indicated by co-localization of RABV P and NEFM (Fig. 4M,N).

### Pronounced infection of various tissues and sensory neurons of the facial head region

Since centrifugal spread to the salivary glands is considered an inevitable step in RABV transmission to other hosts, the extent of RABV infection in tissues of the facial head regions was investigated by imaging of coronal head slices (Fig. S2B). Except for the olfactory bulb, the brain was removed prior to sectioning for separate processing. 3D volume reconstruction from light sheet microscopy z-stacks demonstrated abundant RABV P specific signals throughout various areas of the coronal head slice both after i.c. (Fig. 5A,B; Video S5) and i.m. inoculation (Fig. 6A,B, Fig. S4; Video S6). RABV-infected cells could be identified in the orbital (Fig. 5C,D and 6C,D), nasal (Fig. 5E,F and 6E,F), and oral cavity (Fig. 5G-J and 6G-J). Morphologies of the infected cells revealed a structure typical for sensory neurons in the olfactory epithelium and retina (Fig. 5D,F; Fig. S4) and indicated selective infection of neuronal cells in these tissues. Notably, even though abundant route-independent infection of the olfactory bulb was detectable, presence of fewer RABV-infected cells after i.m. inoculation as compared to the i.c. route demonstrated inoculation route-dependent kinetics of the centrifugal spread to peripheral nerves (Fig. 5A and 6A). Similar to the infection of femoral tissues (Fig. 2 and 3), RABV P was arranged around NEFM-positive axons (Fig.5J, Fig 7A-E), indicating involvement of SCs. Accordingly, RABV was detected at the convex side of the myelin sheaths (Fig. 7G-J, arrowheads; Video S7). This confirmed infection of neuroglia in RABV infection of peripheral nerves of the facial head region. However, in single cases RABV P was also present inside the myelin sheaths, indicating RABV P accumulation in the respective axons (Fig. 7I,J, stars).

**Fig. 5.**
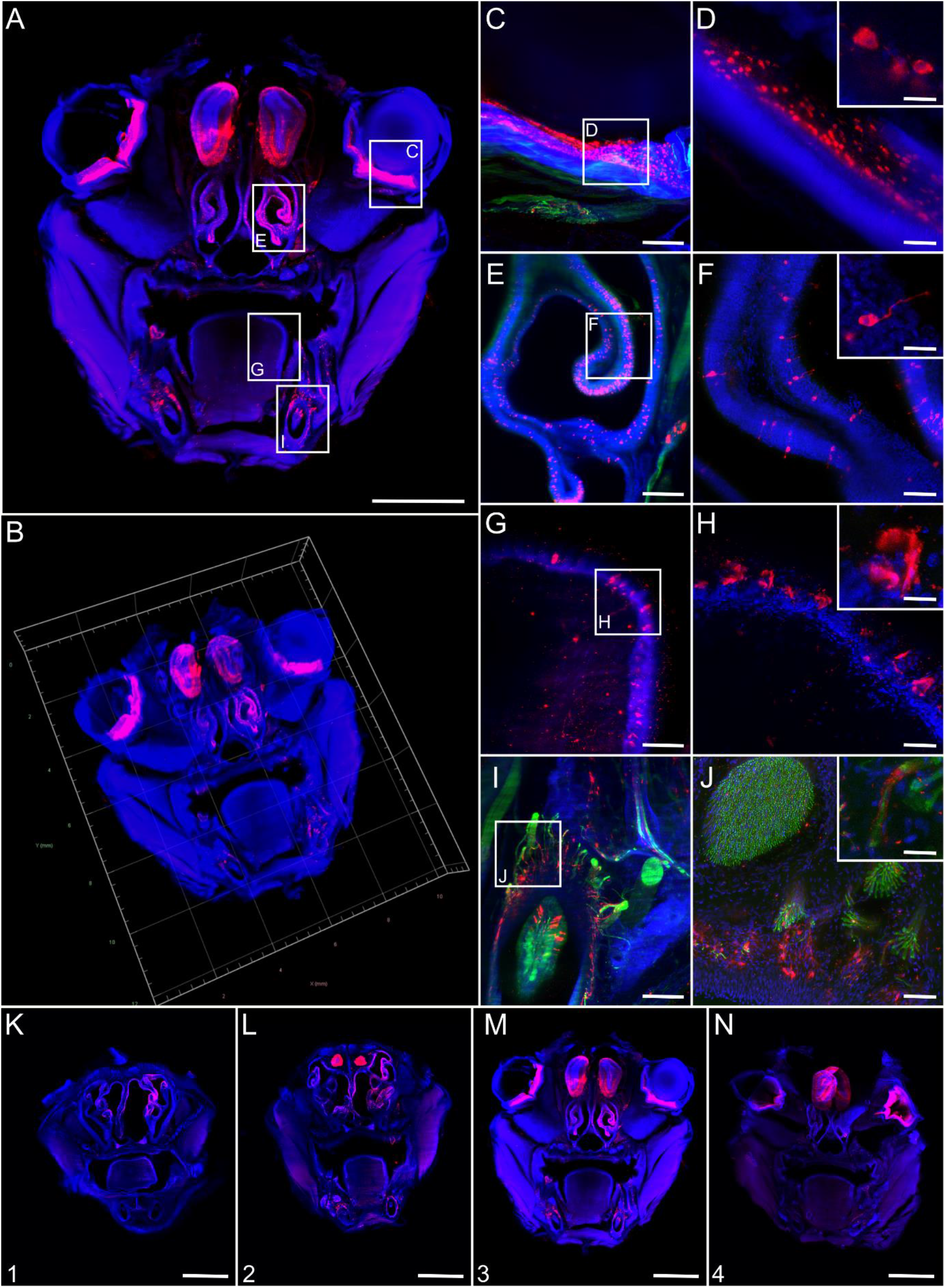
Coronal overview of field RABV-infected mouse head sections after i.c. inoculation. **(A,B)** Maximum z- (A) and 3D projection (B) of mouse head sections after intracranial infection with field RABV [1.26x magnification; z = 1,606 μm]. Indirect immunofluorescence staining against RABV P (red), NEFM (green), and nuclei (blue). Green fluorescence for NEFM not shown as separation from green autofluorescence was not possible at low resolution. Scale bar: 2,000 μm. **(C, E, G, I)** Maximum z-projections of details from A (white boxes) at a magnification of 12.6x. Scale bar: 200 μm. **(D, F, H, J)** Confocal high-resolution z-stacks (corresponding to the indicated regions in C, E, G, I, respectively) [z = 35 μm (D), 32 μm (F), 36 μm (H), 40μm (J); scale bar: 50 μm and 15 μm in detail]. Note: due to the separate confocal image acquisition, the orientations differ from the light sheet microscopy images in C, E, G, and I. **(C,D)** Infected eye, **(E, F)** nasal epithelium, **(G, H)** tongue, **(I, J)** and teeth in mouse head coronal section. **(K-N)** Maximum z-projection of serial mouse head sections (Fig. S2B) [1.26x; z = 1,780 μm (K), 1,820 μm (L), 1,606 μm (M), 1,760 μm (N); green not shown]. Scale bar: 2,000 μm.

**Fig. 6.**
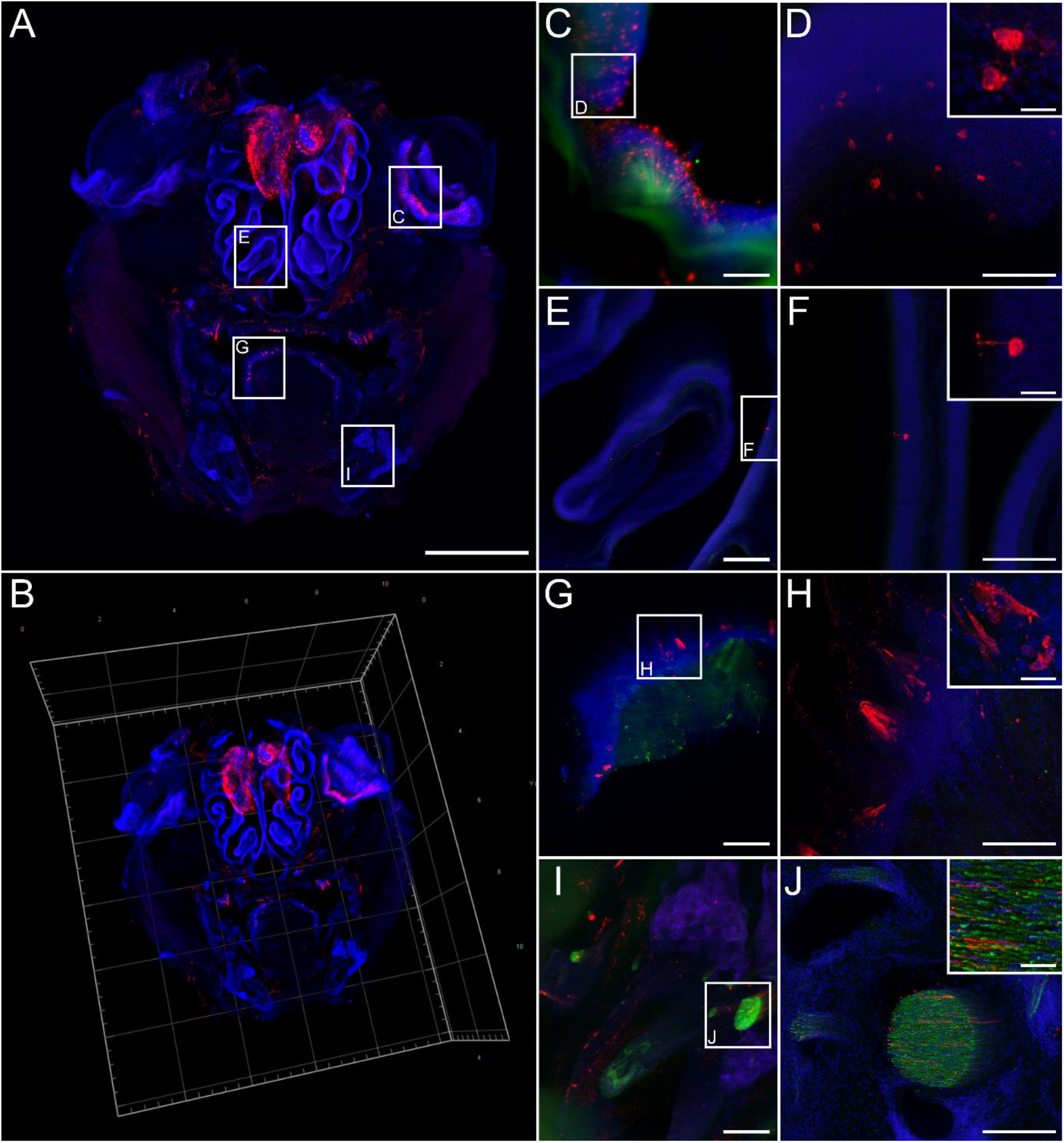
RABV distribution in head cross sections after i.m. inoculation. **(A,B)** Maximum z- (A) and 3D projection of coronal mouse head sections after i.m. infection with field RABV rRABV Dog [1.26x magnification; z = 4,210 μm]. Indirect immunofluorescence staining against RABV P (red), NEFM (green), and nuclei (blue) revealed a markedly reduced degree of infection after i.m. infection in comparison to i.c. infection (see Fig. 6). **(C, E, G, I)** Maximum z-projections of details from A (white boxes) [12.6x; magnification] (Scale bar: 200 μm). **(D, F, H, J)** Confocal high-resolution z-stacks (corresponding to **C, E, G, I**, respectively) [z = 80 μm (D), 26 μm (E), 72 μm (H), 74 μm (J); scale bar: 100 μm] with details (scale bar: 15 μm). **(C,D)** infected eye, **(E, F)** nasal epithelium, **(G, H)** tongue, **(I, J)** and teeth in mouse head coronal section.

**Fig 7.**
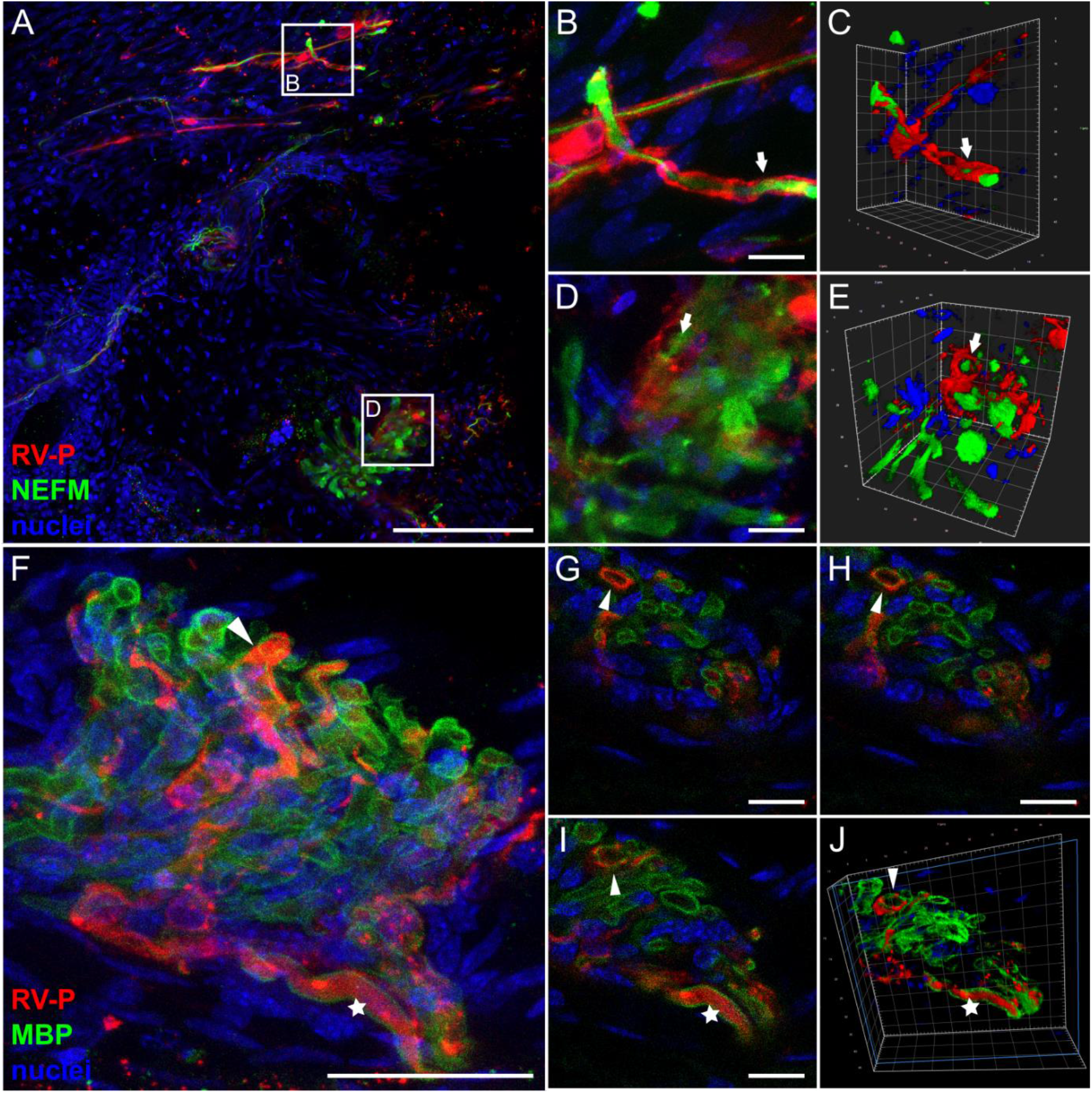
RABV P in axons and infected Schwann cells in coronal head sections. **(A)** Maximum z-projection of high-resolution confocal z-stacks of mouse head sections after i.c. inoculation and staining for RABV P (red), NEFM (green) and nuclei (blue) [z = 38 μm]. Arrows indicate RABV P surrounding NEFM. Scale bar: 100 μm. **(B-E)** Maximum z-projections (B, D) and corresponding 3D projections (C, E) of details of (A) (white boxes). **(F)** Maximum z-projection of confocal z-stacks of mouse head sections after i.c. inoculation stained for RABV P (red), MBP (green) and nuclei (blue) [z = 29 μm]. Scale bar: 25 μm. **(G-I)** Single slices of (F). Arrowhead: RABV P surrounding MBP-positive myelin sheath. Star: RABV P surrounded by MBP-positive myelin sheath. Scale bar: 5 μm. **(J)** 3D projection of (G-H).

## Discussion

Using organic solvent-based organ clearing [45,49] for the 3D deep tissue imaging of RABV infections [69], we aimed at investigating and systematically analyzing the *in vivo* cycle and main checkpoints of a highly virulent field RABV (rRABV Dog) on its journey through the host’s body. To this end, we provide an unprecedented comprehensive overview of the local field RABV tissue tropism and spread in the late clinical phase of RABV encephalitis based on high-volume immunofluorescence imaging. By analyzing peripheral tissues after both i.m. and i.c. infection, we were not only able to visualize RABV infection in PNS and CNS neurons after centripetal and centrifugal spread but also provide clear evidence for the infection of non-neuronal myelinating SCs in peripheral nerves of hind legs and facial head regions (Fig. 2, 3 and 7). SCs constitute a glial cell type in the PNS that functionally correlates to myelinating oligodendrocytes in the CNS (reviewed in [20]). Another important finding is the corroboration of anterograde transport and spread of RABV in axons of peripheral nerves by infection of the SCs after centrifugal spread from the CNS.

Myelinating SCs are peripheral neuroglia, which form the myelin sheath around axons of motor and sensory neurons and, like oligodendrocytes in the CNS, are involved in saltatory conduction and trophic support for neurons (reviewed in [51]). In peripheral nerves, RABV infection was associated with axon demyelination [37,38,57,14,55]. T-lymphocyte-dependent immune pathogenesis was identified to be causative for RABV-mediated neurotic paralysis, the disintegration of myelin sheaths, and the degeneration of axons [65]. Furthermore, there has been speculation that involvement of rabies-induced neuropathogenesis in peripheral nerves results in the typical rabies sings of muscle spasms and deglutition [37].

Although there are indications that SCs are somehow affected by RABV infections [14,37,38,55,57], evidence for direct RABV infection of SCs in the PNS is lacking. While released RABV particles were detected at nodes of Ranvier (known as myelin-sheath gaps occurring along a myelinated axon [40]) and between axonal and SC plasma membranes [31,41,42]) only once infection of SCs is described [4]. Others failed to detect RABV infected SCs (reviewed in [50]). Possible explanations for that may be the use of ultrathin sections in electron microscopy in previous studies that run the inherent risk of missing rare events, RABV strain-specific differences in accumulation in peripheral axons and SCs, and variations in the clinical phase of the analyzed animals.

SCs are immunocompetent and exert key functions in immune regulation by antigen presentation, pathogen detection, and cytokine production [24,68]. Accordingly, productive or non-productive infection of these cells could be critically important to immediate innate responses, local inflammatory processes, and the development of adaptive immunity. Recently, we demonstrated that field RABVs, including the recombinant clone rRABV Dog used in this study, are able to infect astroglia in the CNS to a remarkable degree comparable to neuron infections. Whereas infection of astrocytes seems to be a hallmark of field RABVs, more attenuated lab strains are characterized by highly restricted or absent infection of astrocytes. Accordingly, establishment of astroglia infection by field RABVs was hypothesized to limit or block respective immune responses, subsequent virus elimination, and delay immune pathology [48]. RABV infection of SCs in the PNS, as demonstrated in our study (Figs. 2, 3 and 7), seems to represent a direct correlate to astroglia infection in the CNS. Comparable to the latter, field RABV replication in peripheral neuroglia may result in local suppression of glia-mediated innate immune responses and, thus, gain ability to replicate in SCs. In contrast, similar to the astroglia infection in the brain [46,48], less virulent RABVs may infect SCs in an abortive manner because of a strong type I interferon response. Evasion of innate and intrinsic antiviral pathways have been attributed to the RABV nucleoprotein (N), P, and matrix (M) proteins [10,9,8,36,62]. As discussed above, there is reason to believe that, similar to current hypotheses about the role of neuroglia infection in the CNS, virus variants are not or only inefficiently able to establish infection in peripheral neuroglia. This may induce stronger local interferon responses, more severe local immune pathogenesis, reduced virus replication, and damage of infected neurons. While we could even confirm infection of SCs by another, bat-associated field RABV (Fig. S5), analyses with less virulent lab-adapted strains and live-attenuated vaccine viruses have to be subject of further studies to assess whether the virulence-dependent difference in glial cell infection observed in the brain [48] can be transferred to the SC infections in the PNS described in this study.

Virus infection of peripheral SCs in hind legs and facial head regions by centrifugal spread after i.c. inoculation (Figs. 3 and 7) as well as infection of sensory neurons in the olfactory epithelium and other tissues (Figs. 5 and 6) provide evidence for anterograde transport of RABV along axons *in vivo*, something that has been attributed to late phases of infection [16]. The long distance from the neuron soma in the respective ganglia and the tight spatial association of the infected neuroglia with the axons makes it exceedingly unlikely that virus release at cell bodies and extra-neuronal spread is responsible for distal SC infections.

Both *in vitro* and *in vivo* virus tracking already demonstrated that anterograde RABV transport through axons of peripheral DRG neurons and other peripheral neurons is possible [3,60,6]. However, anterograde virus spread was discussed to occur only in a passive, kinesin-independent manner and the release of infectious virus was called into question [61]. Fast, post-replicative, and glycoprotein (G)-dependent anterograde axonal transport of RABV in cultured DRG neurons, however, provided evidence for active transport to distal axon sites [6]. One reason for the discrepancy observed between *in vitro* tracking and *in vivo* trans-neuronal tracing approaches could be the role of peripheral neuroglia as major innate immune mediators in infected nerves as discussed earlier. Since the standard neuronal tracer virus CVS-11 is less effective in infecting astrocytes in the brain [48], a strong local antiviral immune response at peripheral sites may hinder CVS-11 replication in neuroglia target cells. Moreover, anterogradely transported virus particles in axons may remain below the limit of detection in conventional *in vivo* tracing approaches. Accordingly, anterograde transport in CVS-11 infection may have been overlooked because peripheral neuroglia, which were used here as an indicator for anterograde spread of the used field RABV, might not or only get abortively infected.

Beyond RABV spread in peripheral hind leg nerves after i.m. and i.c. infection, a more generalized roadmap of RABV infection in the CNS and PNS could be build. As for human [58,32] and numerous animal samples (e.g. [34,29,28]), abundant RABV specific signals in the spinal cord and brain could be observed (Fig. 4 and Fig. S6). Virus detection in the orbital, nasal, and oral cavity confirmed previous reports about presence of RABV and related lyssaviruses in a variety of other tissues [17,30,22,1], including retinal ganglion cells, their axons, and lacrimal glands [25,15]. Considering the latter, a potential role of lacrimal glands in virus secretion has been discussed [15]. Whereas lacrimal fluid may not contribute to the transmission of infectious virus to new hosts via saliva, multiple infected sensory neurons in the olfactory and tongue epithelia (Fig. 5 and 6) with a direct connection to the nasopharynx raise the question whether virus secretion at these sites could contribute to the total amount of infectious virus in the salvia. For bat lyssaviruses, for instance, viral antigen was also detected in taste buds of experimentally [22] and naturally infected animals [52].

## Conclusions

By providing a comprehensive and detailed overview of RABV distribution in peripheral tissues after natural (i.m.) and artificial (i.c.) routes of inoculation, we highlight novel insights in the cell tropism and *in vivo* spread of field RABV. Importantly, by unveiling the infection of peripheral neuroglia, we suggest a model in which field RABV infection of immunocompetent SCs is critically important for local innate immune responses in peripheral nerves. This may not only contribute to the delayed immune detection of field RABVs through RABV mediated innate immune response suppression in field virus infected SCs, but may also may explain the high neuroinvasive potential of field RABVs when compared with less virulent lab virus strains, and manifestation of clinical symptoms observed in infected animals and humans. Moreover, by demonstrating infection of myelinating SCs after centrifugal spread from infected brains, we provide evidence for anterograde spread of infectious RABV through axons of the peripheral nerve systems in clinical phases of rabies encephalitis.

## Abbrevitations

BA: benzyl alcohol
BB: benzyl benzoate
CNS: central nervous system
DPE: diphenyl ether
DRG: dorsal root ganglion
EDTA: ethylenediaminetetraacetic acid
G: glycoprotein
i.m.: intramuscular
i.c.: intracranial
LALFF M-V: State Office for Agriculture, Food Safety, and Fishery in Mecklenburg-Western Pomerania
M: matrix protein
MBP: marker myelin basic protein
N: nucleoprotein
NEFM: neurofilamten M
P: phosphoprotein
PNS: peripheral nervous system
RABV: rabies virus
PFA: paraformaldehyde
RNPs: ribonucleoproteins
SC: Schwann cell
TBA: *tert*-butanol

## Declarations

### Ethics Approval

No human material was used in this study. Animal trials have been evaluated by the ethics committee of the State Office for Agriculture, Food Safety, and Fishery in Mecklenburg-Western Pomerania (LALFF M-V) and gained approval with permission 7221.3-2-047/19 and 7221.3-2.1-001/18.

## Acknowledgements

We thank Angela Hillner and Katrin Giesow for technical assistance.

## Supplemantary Material

Supplementary Figures_S1-S6.pdf (including figure captions and descriptions)

Captions Videos S1-S7 (Captions and descriptions for supplementary videos)

Video S1.mp4

Video S2 reduced.avi

Video S3.mp4

Video S4.mp4

Video S5.mp4

Video S6.mp4

Video S7.avi

## Notes

### Competing Interest Statement

The authors have declared no competing interest.

## References

1. Allendorf SD, Cortez A, Heinemann MB, Harary CM, Antunes JM, Peres MG et al (2012) Rabies virus distribution in tissues and molecular characterization of strains from naturally infected non-hematophagous bats. Virus research 165:119–125. doi:10.1016/j.virusres.2012.01.011

2. Amarasinghe GK, Arechiga Ceballos NG, Banyard AC, Basler CF, Bavari S, Bennett AJ et al (2018) Taxonomy of the order Mononegavirales: update 2018. Archives of virology 163:2283–2294. doi:10.1007/s00705-018-3814-x

3. Astic L, Saucier D, Coulon P, Lafay F, Flamand A (1993) The CVS strain of rabies virus as transneuronal tracer in the olfactory system of mice. Brain Res 619:146–156. doi:10.1016/0006-8993(93)91606-s

4. Atanasiu P, Sisman J (1967) [Morphological aspects of rabies virus]. Bull Off Int Epizoot 67:521–533

5. Balachandran A, Charlton K (1994) Experimental rabies infection of non-nervous tissues in skunks (Mephitis mephitis) and foxes (Vulpes vulpes). Vet Pathol 31:93–102. doi:10.1177/030098589403100112

6. Bauer A, Nolden T, Schroter J, Romer-Oberdorfer A, Gluska S, Perlson E et al (2014) Anterograde glycoprotein-dependent transport of newly generated rabies virus in dorsal root ganglion neurons. Journal of virology 88:14172–14183. doi:10.1128/JVI.02254-14

7. Begeman L, GeurtsvanKessel C, Finke S, Freuling CM, Koopmans M, Muller T et al (2018) Comparative pathogenesis of rabies in bats and carnivores, and implications for spillover to humans. Lancet Infect Dis 18:e147–e159. doi:10.1016/S1473-3099(17)30574-1

8. Besson B, Sonthonnax F, Duchateau M, Ben Khalifa Y, Larrous F, Eun H et al (2017) Regulation of NF-kappaB by the p105-ABIN2-TPL2 complex and RelAp43 during rabies virus infection. PLoS Pathog 13:e1006697. doi:10.1371/journal.ppat.1006697

9. Brzozka K, Finke S, Conzelmann KK (2005) Identification of the rabies virus alpha/beta interferon antagonist: phosphoprotein P interferes with phosphorylation of interferon regulatory factor 3. Journal of virology 79:7673–7681. doi:10.1128/JVI.79.12.7673-7681.2005

10. Brzozka K, Finke S, Conzelmann KK (2006) Inhibition of interferon signaling by rabies virus phosphoprotein P: activation-dependent binding of STAT1 and STAT2. Journal of virology 80:2675–2683. doi:10.1128/JVI.80.6.2675-2683.2006

11. Casals J (1940) Influence of Age Factors on Susceptibility of Mice to Rabies Virus. J Exp Med 72:445–451. doi:10.1084/jem.72.4.445

12. Charlton KM, Casey GA (1979) Experimental rabies in skunks: immunofluorescence light and electron microscopic studies. Lab Invest 41:36–44

13. Charlton KM, Nadin-Davis S, Casey GA, Wandeler AI (1997) The long incubation period in rabies: delayed progression of infection in muscle at the site of exposure. Acta Neuropathol 94:73–77. doi:10.1007/s004010050674

14. Chopra JS, Banerjee AK, Murthy JM, Pal SR (1980) Paralytic rabies: a clinico-pathological study. Brain 103:789–802. doi:10.1093/brain/103.4.789

15. Dalton MF, Siepker CL, Maboni G, Sanchez S, Rissi DR (2020) Ocular and Lacrimal Gland Lesions in Naturally Occurring Rabies of Domestic and Wild Mammals. Vet Pathol 57:409–417. doi:10.1177/0300985820911458

16. Davis BM, Rall GF, Schnell MJ (2015) Everything You Always Wanted to Know About Rabies Virus (But Were Afraid to Ask). Annu Rev Virol 2:451–471. doi:10.1146/annurev-virology-100114-055157

17. Fekadu M, Shaddock JH (1984) Peripheral distribution of virus in dogs inoculated with two strains of rabies virus. Am J Vet Res 45:724–729

18. Finke S, Conzelmann KK (2005) Replication strategies of rabies virus. Virus research 111:120–131. doi:10.1016/j.virusres.2005.04.004

19. Fischer M, Freuling CM, Muller T, Wegelt A, Kooi EA, Rasmussen TB et al (2014) Molecular double-check strategy for the identification and characterization of European Lyssaviruses. Journal of virological methods 203:23–32. doi:10.1016/j.jviromet.2014.03.014

20. Fontenas L, Kucenas S (2017) Livin’ On The Edge: glia shape nervous system transition zones. Curr Opin Neurobiol 47:44–51. doi:10.1016/j.conb.2017.09.008

21. Fooks AR, Cliquet F, Finke S, Freuling C, Hemachudha T, Mani RS et al (2017) Rabies. Nat Rev Dis Primers 3:17091. doi:10.1038/nrdp.2017.91

22. Freuling C, Vos A, Johnson N, Kaipf I, Denzinger A, Neubert L et al. (2009) Experimental infection of serotine bats (Eptesicus serotinus) with European bat lyssavirus type 1a. The Journal of general virology 90:2493–2502. doi:10.1099/vir.0.011510-0

23. Gluska S, Zahavi EE, Chein M, Gradus T, Bauer A, Finke S et al (2014) Rabies Virus Hijacks and accelerates the p75NTR retrograde axonal transport machinery. PLoS Pathog 10:e1004348. doi:10.1371/journal.ppat.1004348

24. Gold R, Archelos JJ, Hartung HP (1999) Mechanisms of immune regulation in the peripheral nervous system. Brain Pathol 9:343–360. doi:10.1111/j.1750-3639.1999.tb00231.x

25. Haltia M, Tarkkanen A, Kivela T (1989) Rabies: ocular pathology. Br J Ophthalmol 73:61–67. doi:10.1136/bjo.73.1.61

26. Hampson K, Coudeville L, Lembo T, Sambo M, Kieffer A, Attlan M et al (2015) Estimating the global burden of endemic canine rabies. PLoS Negl Trop Dis 9:e0003709. doi:10.1371/journal.pntd.0003709

27. Hoffmann B, Freuling CM, Wakeley PR, Rasmussen TB, Leech S, Fooks AR et al (2010) Improved safety for molecular diagnosis of classical rabies viruses by use of a TaqMan real-time reverse transcription-PCR “double check” strategy. J Clin Microbiol 48:3970–3978. doi:10.1128/JCM.00612-10

28. Ito FH, Vasconcellos SA, Erbolato EB, Macruz R, Cortes Jde A (1985) Rabies virus in different segments of brain and spinal cord of naturally and experimentally infected dogs. Int J Zoonoses 12:98–104

29. Jackson AC, Reimer DL (1989) Pathogenesis of experimental rabies in mice: an immunohistochemical study. Acta Neuropathol 78:159–165. doi:10.1007/BF00688204

30. Jackson AC, Ye H, Phelan CC, Ridaura-Sanz C, Zheng Q, Li Z, Wan X et al (1999) Extraneural organ involvement in human rabies. Lab Invest 79:945–951

31. Jenson AB, Rabin ER, Bentinck DC, Melnick JL (1969) Rabiesvirus neuronitis. Journal of virology 3:265–269

32. Juntrakul S, Ruangvejvorachai P, Shuangshoti S, Wacharapluesadee S, Hemachudha T (2005) Mechanisms of escape phenomenon of spinal cord and brainstem in human rabies. BMC Infect Dis 5:104. doi:10.1186/1471-2334-5-104

33. Klingen Y, Conzelmann KK, Finke S (2008) Double-labeled rabies virus: live tracking of enveloped virus transport. Journal of virology 82:237–245. doi:10.1128/JVI.01342-07

34. Kojima D, Park CH, Satoh Y, Inoue S, Noguchi A, Oyamada T (2009) Pathology of the spinal cord of C57BL/6J mice infected with rabies virus (CVS-11 strain). J Vet Med Sci 71:319–324. doi:10.1292/jvms.71.319

35. Li J, McGettigan JP, Faber M, Schnell MJ, Dietzschold B (2008) Infection of monocytes or immature dendritic cells (DCs) with an attenuated rabies virus results in DC maturation and a strong activation of the NFkappaB signaling pathway. Vaccine 26:419–426. doi:10.1016/j.vaccine.2007.10.072

36. Masatani T, Ito N, Shimizu K, Ito Y, Nakagawa K, Sawaki Y et al (2010) Rabies virus nucleoprotein functions to evade activation of the RIG-I-mediated antiviral response. Journal of virology 84:4002–4012. doi:10.1128/JVI.02220-09

37. Minguetti G, Hofmeister RM, Hayashi Y, Montano JA (1997) Ultrastructure of cranial nerves of rats inoculated with rabies virus. Arq Neuropsiquiatr 55:680–686. doi:10.1590/s0004-282x1997000500002

38. Minguetti G, Negrao MM, Hayashi Y, de Freitas OT (1979) Ultrastructure of peripheral nerves of mice inoculated with rabies virus. Arq Neuropsiquiatr 37:105–112. doi:10.1590/s0004-282x1979000200001

39. Morimoto K, Hooper DC, Carbaugh H, Fu ZF, Koprowski H, Dietzschold B (1998) Rabies virus quasispecies: implications for pathogenesis. Proceedings of the National Academy of Sciences of the United States of America 95:3152–3156. doi:10.1073/pnas.95.6.3152

40. Murphy FA (1977) Rabies pathogenesis. Archives of virology 54:279–297. doi:10.1007/BF01314774

41. Murphy FA, Bauer SP, Harrison AK, Winn WC, Jr. (1973) Comparative pathogenesis of rabies and rabies-like viruses. Viral infection and transit from inoculation site to the central nervous system. Lab Invest 28:361–376

42. Murphy FA, Harrison AK, Winn WC, Bauer SP (1973) Comparative pathogenesis of rabies and rabies-like viruses: infection of the central nervous system and centrifugal spread of virus to peripheral tissues. Lab Invest 29:1–16

43. Nolden T, Pfaff F, Nemitz S, Freuling CM, Hoper D, Muller T et al (2016) Reverse genetics in high throughput: rapid generation of complete negative strand RNA virus cDNA clones and recombinant viruses thereof. Sci Rep 6:23887. doi:10.1038/srep23887

44. Orbanz J, Finke S (2010) Generation of recombinant European bat lyssavirus type 1 and inter-genotypic compatibility of lyssavirus genotype 1 and 5 antigenome promoters. Archives of virology 155:1631–1641. doi:10.1007/s00705-010-0743-8

45. Pan C, Cai R, Quacquarelli FP, Ghasemigharagoz A, Lourbopoulos A, Matryba P, Plesnila N, Dichgans M, Hellal F, Erturk A (2016) Shrinkage-mediated imaging of entire organs and organisms using uDISCO. Nature methods 13:859–867. doi:10.1038/nmeth.3964

46. Pfefferkorn C, Kallfass C, Lienenklaus S, Spanier J, Kalinke U, Rieder M et al (2016) Abortively Infected Astrocytes Appear To Represent the Main Source of Interferon Beta in the Virus-Infected Brain. Journal of virology 90:2031–2038. doi:10.1128/JVI.02979-15

47. Piccinotti S, Whelan SP (2016) Rabies Internalizes into Primary Peripheral Neurons via Clathrin Coated Pits and Requires Fusion at the Cell Body. PLoS Pathog 12:e1005753. doi:10.1371/journal.ppat.1005753

48. Potratz M, Zaeck L, Christen M, Te Kamp V, Klein A, Nolden T et al (2020) Astrocyte Infection during Rabies Encephalitis Depends on the Virus Strain and Infection Route as Demonstrated by Novel Quantitative 3D Analysis of Cell Tropism. Cells 9. doi:10.3390/cells9020412

49. Renier N, Wu Z, Simon DJ, Yang J, Ariel P, Tessier-Lavigne M (2014) iDISCO: a simple, rapid method to immunolabel large tissue samples for volume imaging. Cell 159:896–910. doi:10.1016/j.cell.2014.10.010

50. Rossiter JP, Jackson AC (2020) Pathology. In: Rabies scientific basis of the disease and its management. 4th edn. Academic Press, pp 347–371

51. Salzer JL (2015) Schwann cell myelination. Cold Spring Harb Perspect Biol 7:a020529. doi:10.1101/cshperspect.a020529

52. Schatz J, Teifke JP, Mettenleiter TC, Aue A, Stiefel D, Muller T et al (2014) Lyssavirus distribution in naturally infected bats from Germany. Vet Microbiol 169:33–41. doi:10.1016/j.vetmic.2013.12.004

53. Schindelin J, Arganda-Carreras I, Frise E, Kaynig V, Longair M, Pietzsch T et al (2012) Fiji: an open-source platform for biological-image analysis. Nature methods 9:676–682. doi:10.1038/nmeth.2019

54. Schneider CA, Rasband WS, Eliceiri KW (2012) NIH Image to ImageJ: 25 years of image analysis. Nature methods 9:671–675

55. Tangchai P, Vejjajiva A (1971) Pathology of the peripheral nervous system in human rabies. A study of nine autopsy cases. Brain 94:299–306. doi:10.1093/brain/94.2.299

56. Te Kamp V, Freuling CM, Vos A, Schuster P, Kaiser C, Ortmann S, Kretzschmar A et al (2020) Responsiveness of various reservoir species to oral rabies vaccination correlates with differences in vaccine uptake of mucosa associated lymphoid tissues. Sci Rep 10:2919. doi:10.1038/s41598-020-59719-4

57. Teixeira F, Aranda FJ, Castillo S, Perez M, Del Peon L, Hernandez O (1986) Experimental rabies: ultrastructural quantitative analysis of the changes in the sciatic nerve. Exp Mol Pathol 45:287–293. doi:10.1016/0014-4800(86)90017-1

58. Tirawatnpong S, Hemachudha T, Manutsathit S, Shuangshoti S, Phanthumchinda K, Phanuphak P (1989) Regional distribution of rabies viral antigen in central nervous system of human encephalitic and paralytic rabies. J Neurol Sci 92:91–99. doi:10.1016/0022-510x(89)90178-0

59. Tsiang H (1979) Evidence for an intraaxonal transport of fixed and street rabies virus. J Neuropathol Exp Neurol 38:286–299. doi:10.1097/00005072-197905000-00008

60. Tsiang H, Lycke E, Ceccaldi PE, Ermine A, Hirardot X (1989) The anterograde transport of rabies virus in rat sensory dorsal root ganglia neurons. The Journal of general virology 70 (Pt 8):2075–2085. doi:10.1099/0022-1317-70-8-2075

61. Ugolini G (2010) Advances in viral transneuronal tracing. J Neurosci Methods 194:2–20. doi:10.1016/j.jneumeth.2009.12.001

62. Vidy A, Chelbi-Alix M, Blondel D (2005) Rabies virus P protein interacts with STAT1 and inhibits interferon signal transduction pathways. Journal of virology 79:14411–14420. doi:10.1128/JVI.79.22.14411-14420.2005

63. Vos A, Freuling CM, Hundt B, Kaiser C, Nemitz S, Neubert A et al (2017) Oral vaccination of wildlife against rabies: Differences among host species in vaccine uptake efficiency. Vaccine 35:3938–3944. doi:10.1016/j.vaccine.2017.06.022

64. Vos A, Nolden T, Habla C, Finke S, Freuling C, Teifke J et al (2013) Raccoons (Procyon lotor) in Germany as potential reservoir species for Lyssaviruses. Eur J Wildl Res 59:637–643. doi:10.1007/s10344-013-0714-y

65. Weiland F, Cox JH, Meyer S, Dahme E, Reddehase MJ (1992) Rabies virus neuritic paralysis: immunopathogenesis of nonfatal paralytic rabies. Journal of virology 66:5096–5099

66. Wickersham IR, Finke S, Conzelmann KK, Callaway EM (2007) Retrograde neuronal tracing with a deletion-mutant rabies virus. Nature methods 4:47–49. doi:10.1038/nmeth999

67. Wickersham IR, Lyon DC, Barnard RJ, Mori T, Finke S, Conzelmann KK et al (2007) Monosynaptic restriction of transsynaptic tracing from single, genetically targeted neurons. Neuron 53:639–647. doi:10.1016/j.neuron.2007.01.033

68. Ydens E, Lornet G, Smits V, Goethals S, Timmerman V, Janssens S (2013) The neuroinflammatory role of Schwann cells in disease. Neurobiol Dis 55:95–103. doi:10.1016/j.nbd.2013.03.005

69. Zaeck L, Potratz M, Freuling CM, Muller T, Finke S (2019) High-Resolution 3D Imaging of Rabies Virus Infection in Solvent-Cleared Brain Tissue. Jove-J Vis Exp. doi:ARTN e59402 10.3791/59402

70. Zampieri N, Jessell TM, Murray AJ (2014) Mapping sensory circuits by anterograde transsynaptic transfer of recombinant rabies virus. Neuron 81:766–778. doi:10.1016/j.neuron.2013.12.033

