## Supplementary material for "Neuroglia Infection by Rabies Virus after Anterograde Virus Spread in Peripheral Neurons": Captions Videos S1 to S7

### Captions Videos S1-S7

**Video S1:** **Field RABV-infected nerve fibers in mouse hind leg after i.m. inoculation.** 3D volume reconstruction of RABV P distribution in femoral tissue (see Fig. 1C in manuscript). 12.6x magnification; z = 574 µm. Red: RABV P; blue: nuclei.

**Video S2:** **Detection of RABV P specific signals around tubular NEFM signals.** Tomogram of high-resolution confocal z-stack of RABV-infected nerve fibers. RABV P is not co-localizing with NEFM in axons (arrowheads) but surrounds NEFM positive axons and cells nuclei (arrow). Z = 50 µm. Red: RABV P; green: NEFM; blue: nuclei.

**Video S3:** **RABV P accumulates in the cytomplasm of myelinating Schwann cells.** Volumetric 3D projection of infected nerves in hind leg tissue after i.m. inoculation of field RABV. Staining with Schwann cell marker MBP reveals hollow tubules that are surrounded by RABV P. Cell nuclei in the RABV P positive cells further supports of RABV infection of axon ensheating SCs. Red: RABV P; green: MBP; blue: nuclei.

**Video S4:** **Infection of Schwann cells in hind leg nerve after i.c. inoculation.** Volumetric 3D projection of RABV P distribution in femoral tissue. Three-layered structure with RABV P specific signals around axons ensheated by MBP. Red: RABV P; green: MBP; blue: NEFM.

**Video S5:** **RABV infection of various sites in mouse head cross section.** 3D volume projection from light sheet microscopy z-stack of field RABV infected mouse head after i.c. inoculation. 1.26x magnification, z = 1,606 µm. Red: RABVP; blue: nuclei.

**Video S6:** **Coronal overview of field RABV infected mouse head section after i.m. inoculation.** 3D volume projection from light sheet microscopy z-stack. 1.26x magnification, z = 4,210 µm. Red: RABV P; blue: nuclei.

**Video S7:** **Infected Schwann cells also in facial head region.** Tomogram of high-resolution confocal z-stack of RABV-infected nerve fibers in mouse head after i.c. inoculation. Detection of RABV P at the convex side of the peripheral nerves in mouse head sections (arrowhead), Partially RABV P is also present inside of myelin sheaths, indication also RABV P accumulation in the respective axons (arrow). Z = 16 µm. Red: RABV P; green: MBP; blue: nuclei.
