## Supplementary Figures S1 to S6 for "Neuroglia Infection by Rabies Virus after Anterograde Virus Spread in Peripheral Neurons"

#### Figures S1-S6

### Neuroglia Infection by RABV after Anterograde Virus Spread in Peripheral Neurons

Madlin Potratz, Luca M. Zaeck, Carlotta Weigel, Antonia Klein, Conrad M. Freuling, Thomas Müller, Stefan Finke

*Friedrich-Loeffler-Institut (FLI), Federal Research Institute for Animal Health, Institute of Molecular Virology and Cell Biology, 17493 Greifswald-Insel Riems, Germany*

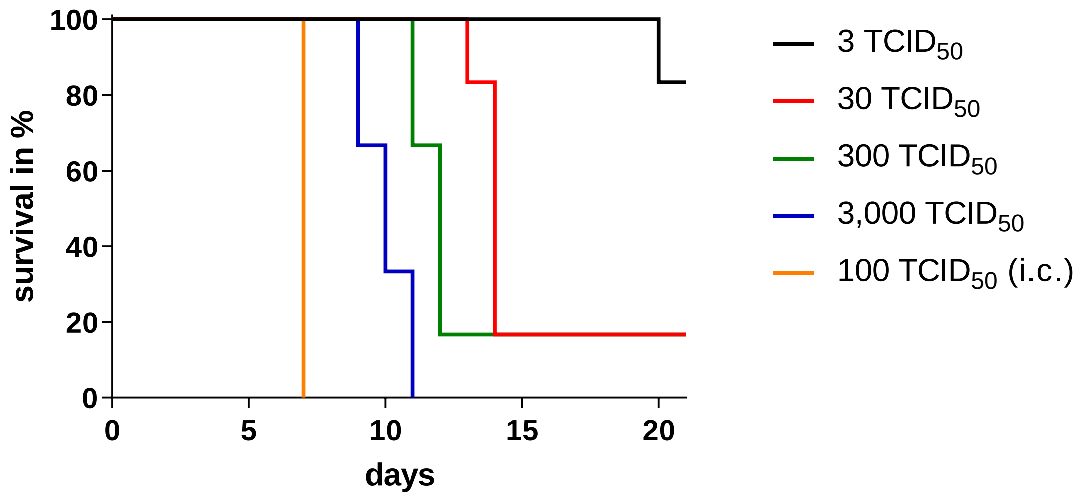


**Fig. S1: Kaplan-Meyer survival plot of RABV infection experiment.** Comparison of dose-dependent survival of mice (six per group) after i.m. inoculation of rRABV Dog (3 to 3,000 $TCID_{50}$). A control group of three mice i.c.-inoculated with 100 $TCID_{50}$ rRABV Dog is shown in orange.

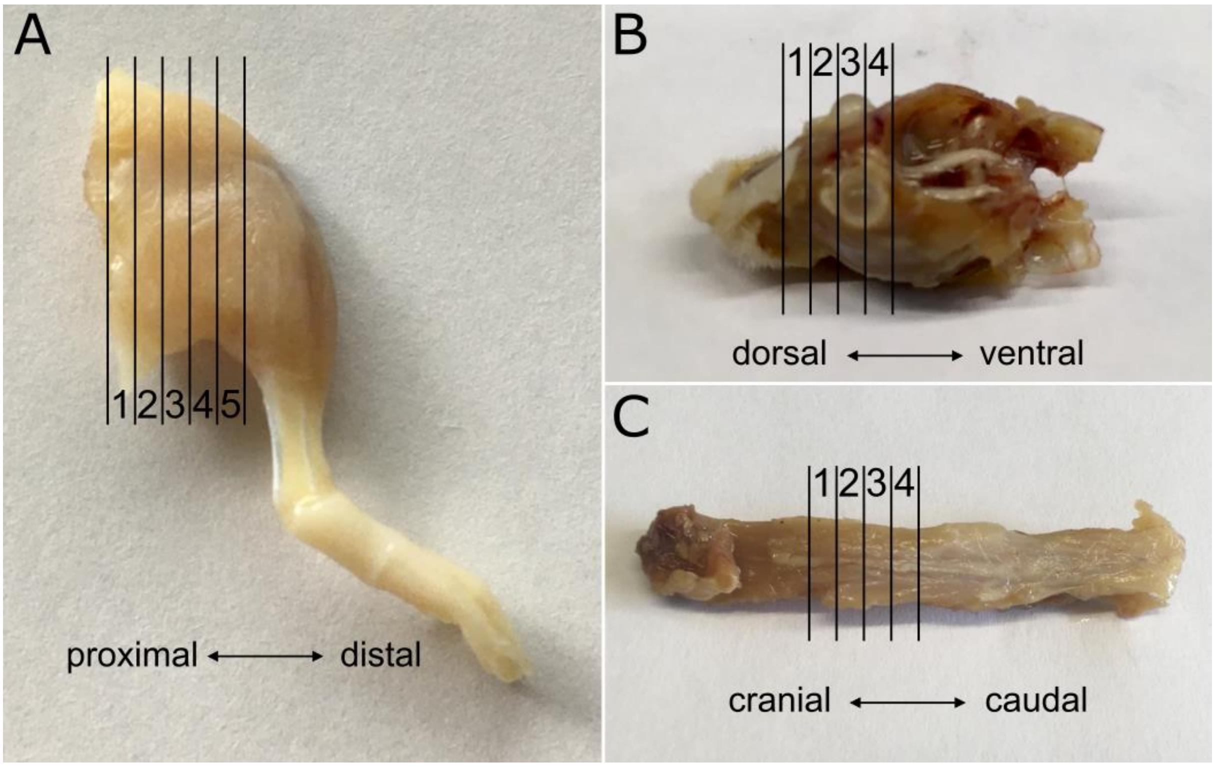


**Fig. S2: Cutting pattern of cross-sections from peripheral tissues of infected mice.** Hind legs (A), heads (B) and spinal columns (C) were decalcified and sectioned with a scalpel into several 1-2 mm thick slices.

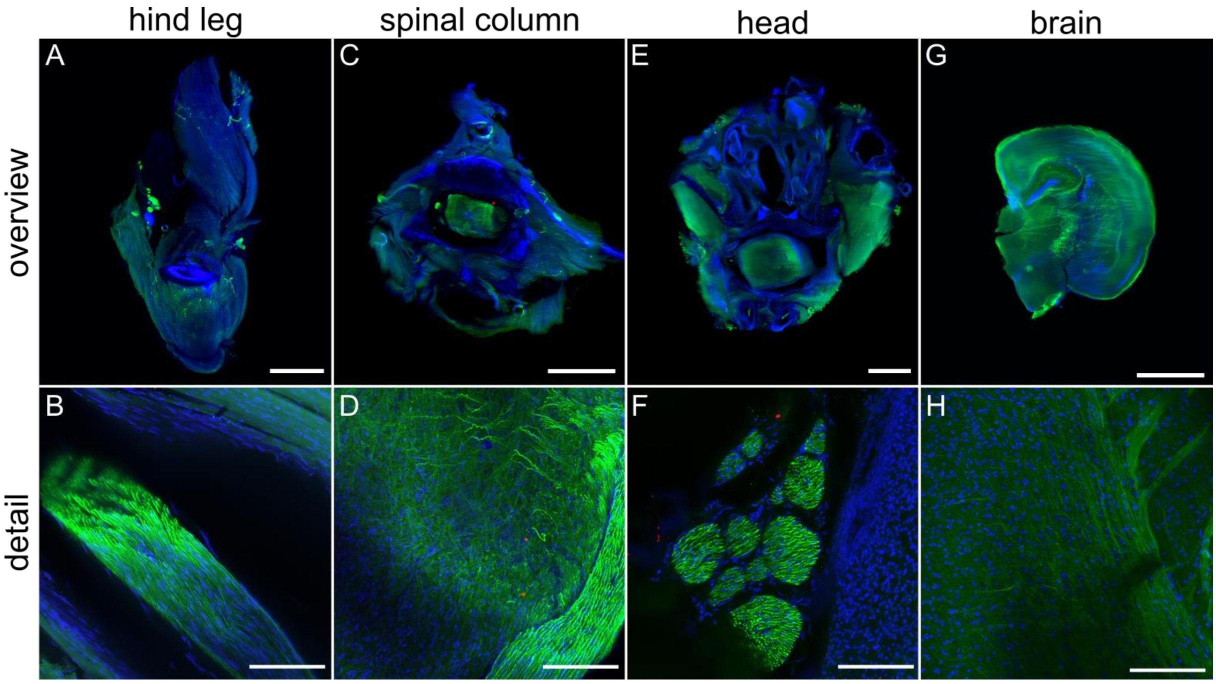


**Fig. S3: Non-infected peripheral mouse tissues after indirect immunofluorescence staining against RV-P (red), NEFM (green) and nuclei (blue).** Maximum z-projections of light sheet overviews and high-resolution confocal z-stacks from hind leg (A,B), spinal column (C,D), head (E,F) and brain (G,H). No RABV antigen was detected. Scale bar: 1,500 µm (overview), 100 µm (detail).

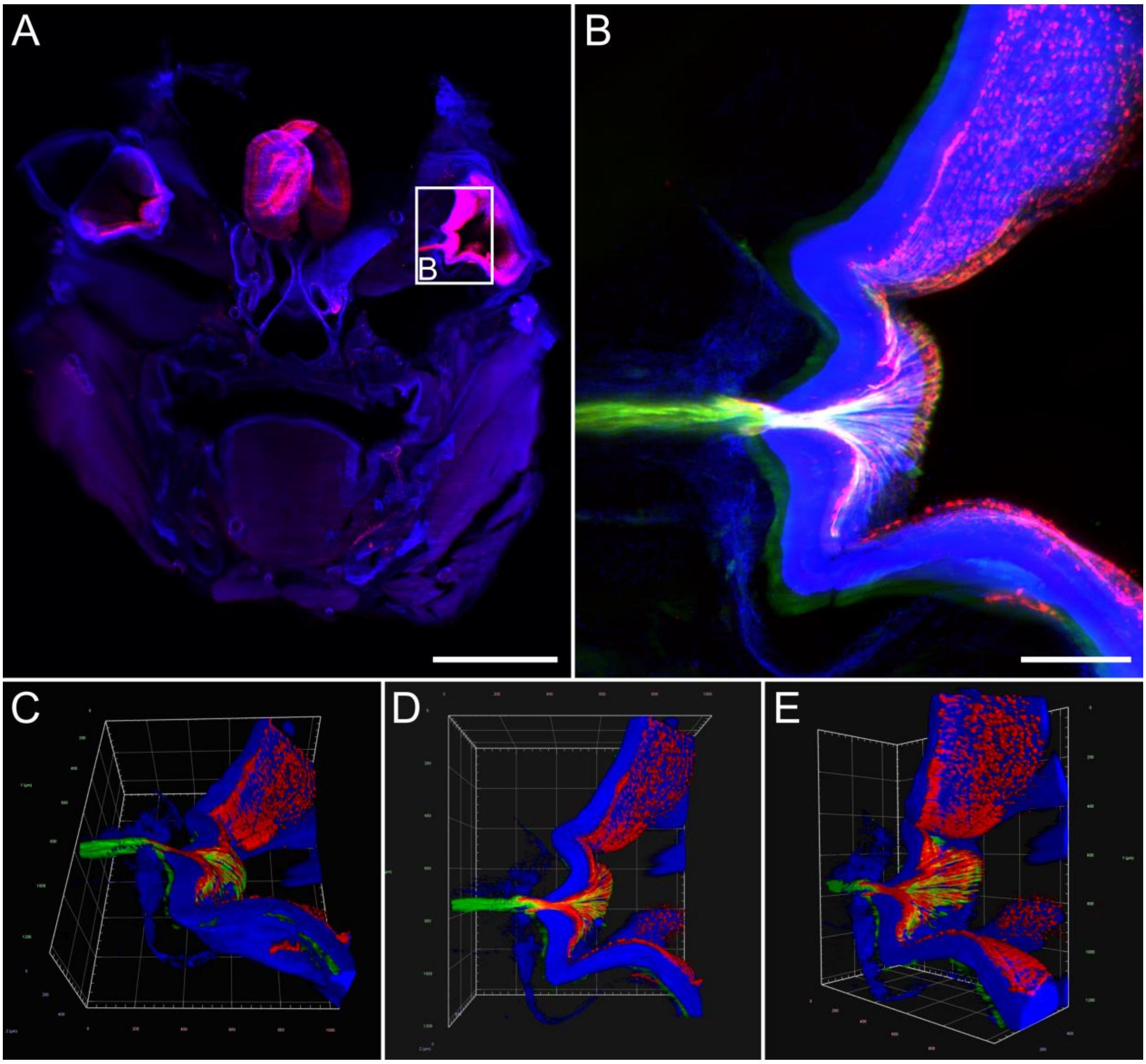


**Fig. S4: Field RABV infection of optic nerve and retina. (A)** Maximum z-projection of coronal mouse head section after i.c. inoculation [1.26x magnification, z = 1,760 µm, Scale bar 2,000 µm]. RV-P (red), nuclei (blue). **(B)** Maximum z-projection of detail from (A) (see white box) with a magnification of 12.6x. Green: NEFM. Scale bar: 200 µm. **(C-E)** Respective 3D projections of (B). Different viewing angles of the infection of the orbital cavity, including the optic nerve, are shown.

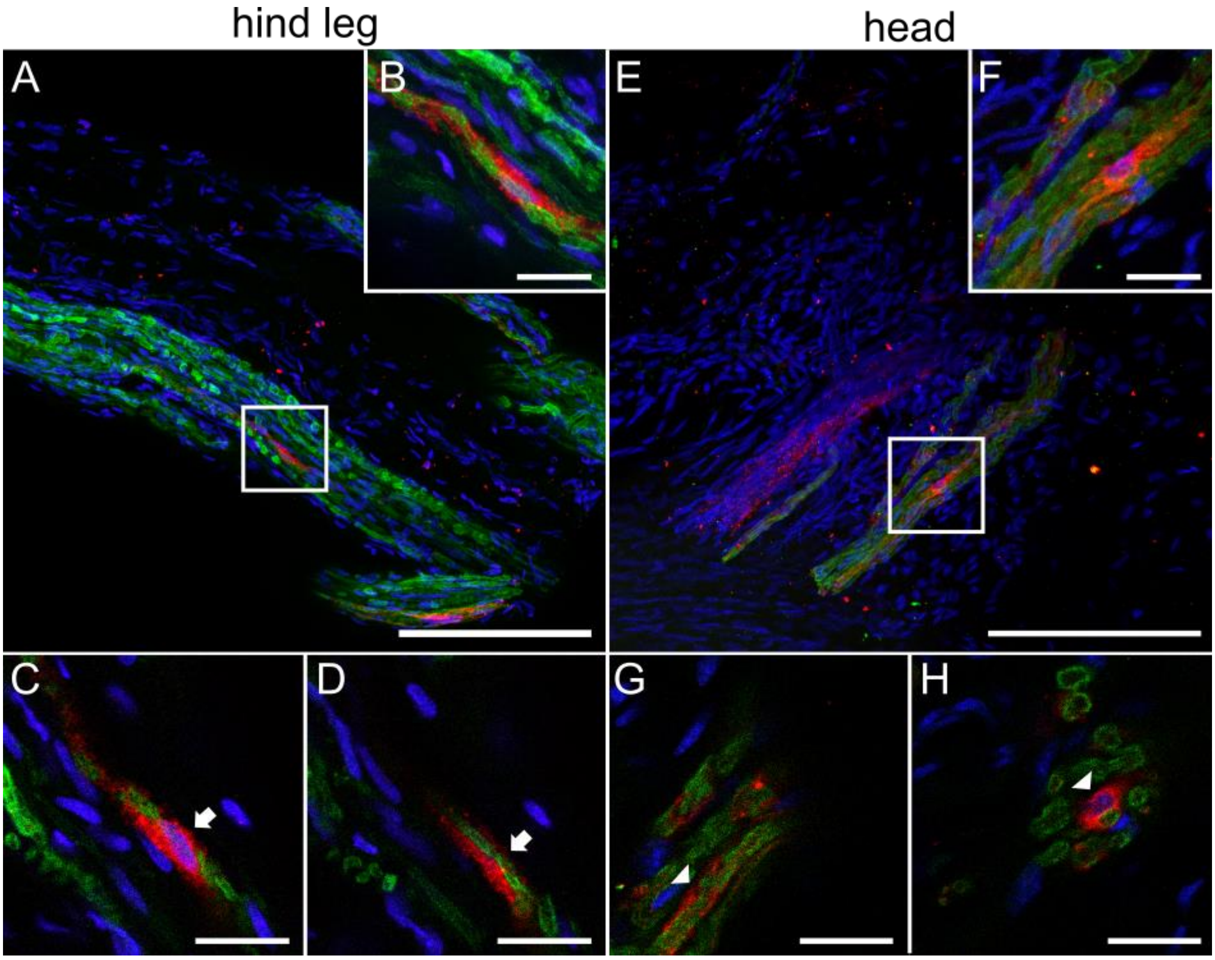


**Fig. S5: A different, bat-associated field RABV demonstrates a comparable pattern of RABV P in hind leg and head nerves, including infected Schwann cells, after i.m. inoculation. (A,B)** Maximum z-projection (A) and detail (B) of confocal high-resolution z-stacks of hind leg section from an i.m.-infected mice with RABV Bat [z = 55 µm; Scale bar: 100 µm (A), 15 µm (B)]. Indirect immunofluorescence staining against RABV P (red), MBP (green), and nuclei (blue). Individual infected nerve fibers were detected, in which RABV P surrounds the MBP signals. **(C,D)** Single planes of detail view from (B). RABV P was detected around MBP signals (white arrows). Scale bar: 15 µm. **(E,F)** Maximum z-projection of nerve fibers in coronal head sections after i.m. inoculation of RABV Bat [z = 31 µm; Scale bar: 100 µm (A), 15 µm (B)]. Indirect immunofluorescence staining against RABV P (red), MBP (green) and nuclei (blue). **(G,H)** Single planes of detail view from (F). RABV P signals were located around MBP structures (white arrowheads). Scale bar: 15 µm.

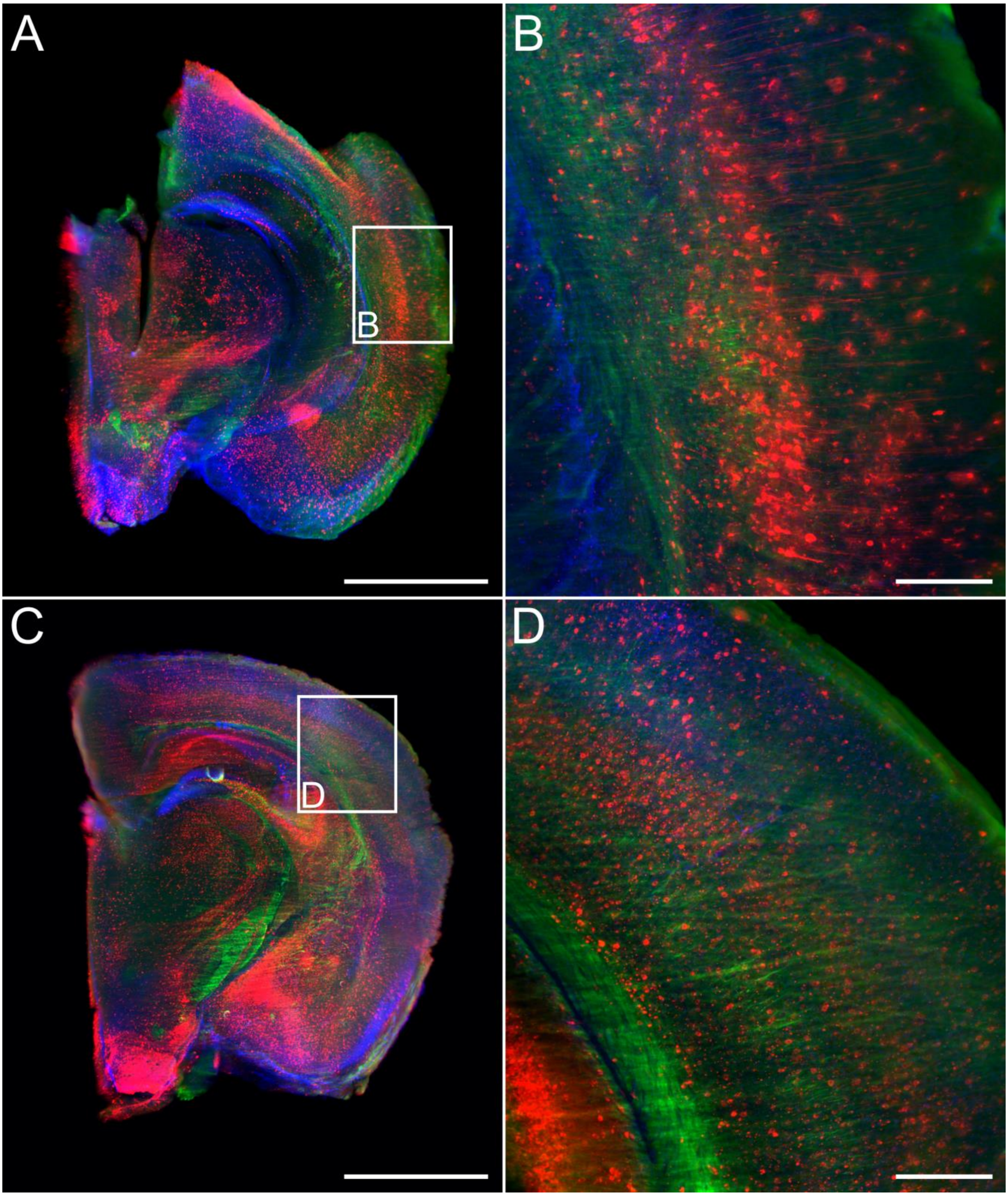


**Fig. S6: Comparison of field RABV-infected brains after i.m. and i.c. inoculation in clinically diseased mice (10 and 7 days post-inoculation, respectively. (A,C)** Independent of the inoculation route, the brains of i.m.- (A) and i.c. (C)-infected mice exhibited massive RABV infection throughout multiple areas of the brain, confirming strong CNS replication independent of the inoculation route. [2.5x magnification; [z = 1,282 µm (A), 1,384 µm (B)]. RABV P = red, NEFM = green, nuclei = blue. Scale bar: 1,500 µm. **(B,D)** Maximum z-projection of details (white boxes) of A and C with magnification of [12.6x;z = 1,130 µm (B), 386 µm (D)]. Scale bar: 200 µm.
